# Sustainable biotransformation of microalgae *via* probiotic fermentation for enhanced functional, nutritional, and sensory properties

**DOI:** 10.1101/2025.08.15.670464

**Authors:** Po-Hsiang Wang, Zann Yi Qi Tan, Choy Eng Nge, Nurhidayah Basri, Lina Xian Yu Lee, Aaron Thong, Mario Wibowo, Elaine Jinfeng Chin, Sharon Crasta, Geraldine Chan, Yoganathan Kanagasundaram, Siew Bee Ng

## Abstract

Microalgae represent a sustainable food source with exceptional CO□ fixation efficiency; however, their integration into the food chain is hindered by undesirable organoleptic properties. This study establishes a green biotransformation platform using **G**enerally **R**ecognized **A**s **S**afe (GRAS) bacterium *Lactiplantibacillus plantarum* to ferment *Chlorella vulgaris* biomass. This fermentation process operates without the use of harsh chemicals and organic solvents, enabling the full utilization of the biomass while improving sensory quality. Notably, the *L. plantarum* fermentation maintained dried biomass weight, in contrast to ∼15–40% loss seen with *Bacillus* spp., further enhancing the carbon-negative profile of microalgae. Tiered olfactory analysis and gas chromatography–mass spectrometry revealed selective reduction of polyunsaturated fatty acid–derived aldehydes and accumulation of flavor-active volatiles, including pyrazines and phenylethyl derivatives. Electronic tongue and liquid chromatography–mass spectrometry confirmed elevated umami taste, *via* increased glutamate and nucleotide levels. Additionally, the fermentation of microalgae with *L. plantarum* converted aromatic amino acids into antioxidant aromatic lactates, exemplifying catalytic, rather than stoichiometric efficiency. Overall, this renewable fermentation strategy converts photoenergy-fuelled, CO□-derived microalgal biomass into direct functional food ingredients under mild, organic solvent-free conditions, while bypassing conventional downstream extraction and purification steps.

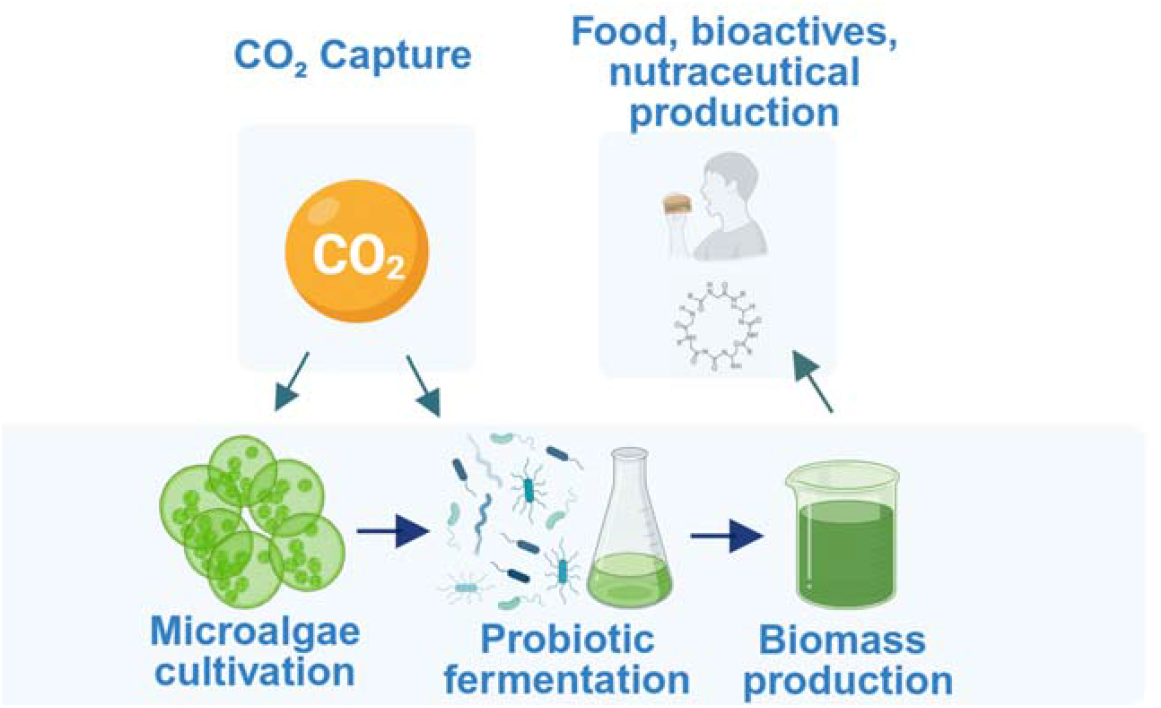

## Introduction

Traditional protein sources and food production methods face significant sustainability challenges, including land usage limitations, excessive water consumption, and substantial carbon footprints ^1^. As such, edible microalgae have emerged as promising candidates for sustainable food production, owing to their exceptional photo-mixotrophic capabilities, high resource efficiency, minimal land requirements, and remarkable carbon capture potential ^2-3^. Historically, microalgae have been extensively utilized in industrial and environmental contexts, including bioremediation of wastewater, CO□ capture, biofuel production, and as sources for commodity biochemicals and high-value compounds ^4-5^. However, their direct application as food ingredients remains comparatively underexplored despite their considerable nutritional value, notably their richness in protein, complex carbohydrates, valuable lipids, and diverse micronutrients. To date, the targeted implementation of microbial fermentation as a method to specifically transform microalgae biomass for direct food production constitutes a significant, yet largely unaddressed, research frontier. The biotransformation of microalgae by carefully selected **G**enerally **R**ecognized **A**s **S**afe (GRAS) probiotic bacteria can enhance nutritional bioavailability and improve sensory characteristics, unlocking tremendous value in alternative protein systems.

Microalgae such as *Chlorella* can fix CO_2_ while utilizing solar energy, with photosynthetic efficiency rates substantially higher than terrestrial crops (10–20% versus 1– 2%) ^6^. Their rapid life cycles, minimal land footprint, and high fertility, coupled with their rich biochemical composition, render them valuable for producing a wide array of chemicals ^7^. Additionally, although differences between and within species exist, microalgae generally produce nutrient-dense biomass containing complete protein profiles and valuable bioactive compounds, such as □-3 polyunsaturated fatty acids (PUFAs), polysaccharides, carotenoids, vitamins, phenolics, and phycobiliproteins ^8^. Despite their nutritional and environmental merits, challenges related to consumer acceptance hinder the integration of microalgae into mainstream food systems. A key factor among these barriers is the undesirable sensory attributes of microalgae, including intense colour and unpleasant flavours which are commonly described as “fishy,” “grassy,” or “pond-like”, significantly impeding palatability ^9^. Additionally, the rigid cell walls of many microalgae species limit nutrient bioavailability and digestibility, further reducing their nutritional efficiency ^10^. These factors have restricted the widespread adoption of microalgae as primary food ingredients despite their exceptional nutritional profiles and environmental credentials.

Biotransformation *via* microbial fermentation presents an elegant green chemistry solution to these challenges. While microalgae-microbial consortia have been explored for industrial applications, their use for direct food enhancement through probiotic fermentation remains underexplored. By employing carefully selected GRAS edible microorganisms as biocatalysts, the biotransformation of microalgae biomass proceeds without requiring harsh chemicals, high temperatures, or energy-intensive processes. Furthermore, microbial metabolism has the potential to modify the negative organoleptic qualities of microalgae while simultaneously enhancing their nutritional aspects by producing additional bioactive compounds. In this study, we investigate the biotransformation of *Chlorella vulgaris* microalgal biomass using selected GRAS probiotic bacteria to address the primary limitations hindering the mainstream adoption of microalgal food through a comprehensive platform approach. We employed an enzymatic-based systematic screening pipeline to identify bacterial strains capable of reducing unpleasant sensory attributes while enhancing functional properties, nutrient bioavailability, and bioactive compound profiles. The selected probiotic strain increased the dried biomass weight by 5%, contrasting with the 15–40% weight reduction observed with *Bacillus* species. Comprehensive analyses using an electronic tongue (E-tongue), olfactory evaluations, liquid chromatography-mass spectrometry (LC-MS) and gas chromatography-mass spectrometry (GC-MS) confirmed the attenuation of unpleasant flavors and the generation of desirable umami taste and aroma compounds. Collectively, these findings establish a foundation for developing microalgae-based functional foods with improved sensory and nutritional qualities via sustainable biotransformation strategies.

## Results and Discussion

### Establishment of a GRAS microbial fermentation platform for microalgae

To unlock the nutritional and sensory potential of microalgae through fermentation, we developed a modular bacterial fermentation platform using food-grade bacterial isolates (**Figure 1**). The workflow commenced by inoculating microalgae with preselected GRAS strains from the **A**gency for **S**cience, **T**echnology, and **R**esearch’s **N**atural **P**roduct **L**ibrary (A*STAR NPL), a comprehensive resource comprising over 37,000 plant specimens and 123,000 microbial strains ^11^. Fermentations were conducted under submerged conditions at 37°C, followed by freeze-drying and sample extraction for comprehensive chemical and sensory profiling. Analytical outputs were generated using a combination of colorimetric enzyme activity assays, olfactory evaluations, E-tongue measurements, GC-MS and LC-MS.

**Figure 1.**
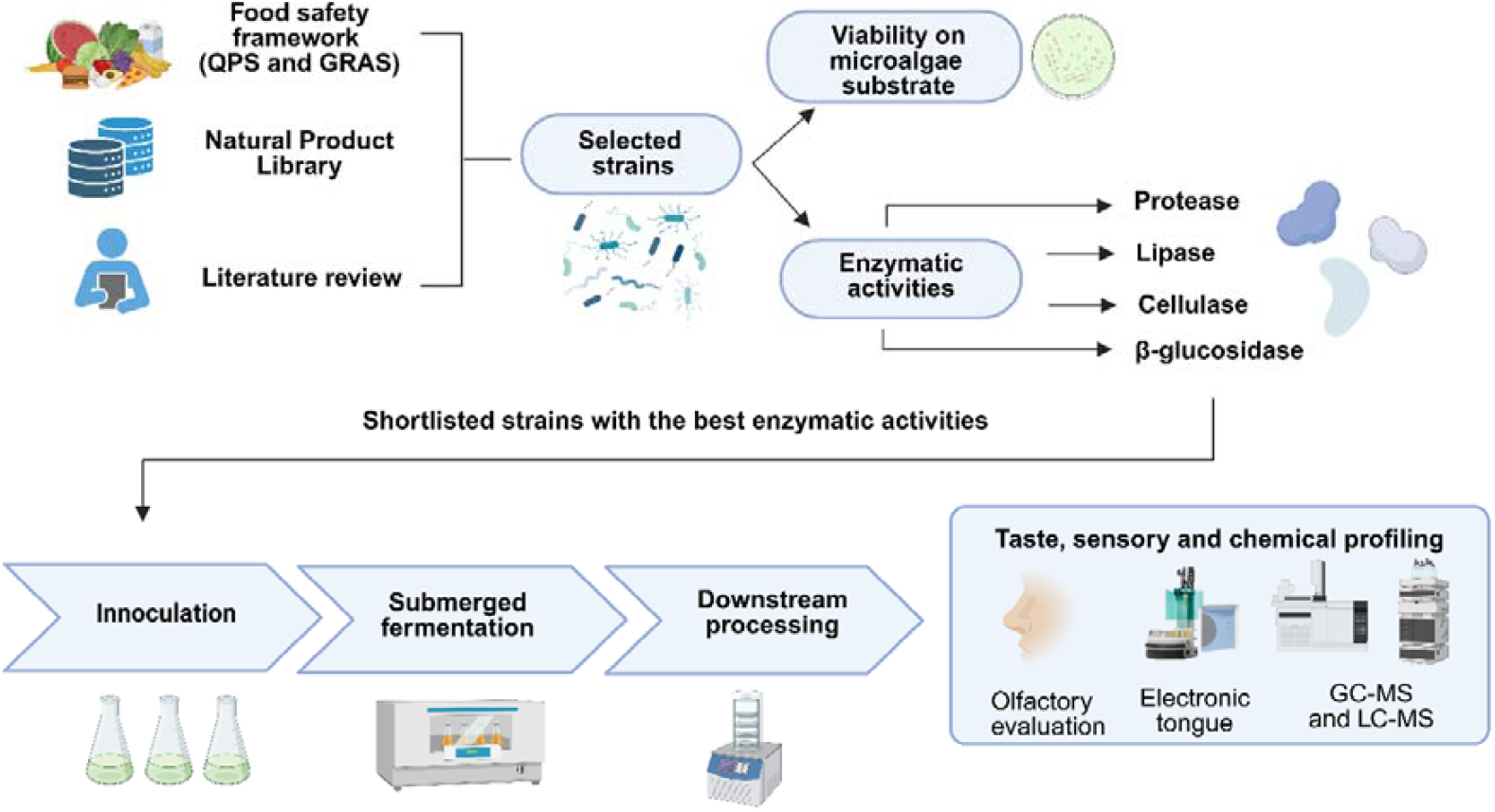
Integrated biotransformation platform showcasing the pipeline for food-grade strain selection, microalgae fermentation, and multi-modal sensory and chemical profiling.

This integrated approach enabled rapid dereplication and high-resolution phenotyping of bacterial fermentates, facilitating the identification of strains and resulting fermentates with desirable nutritional and sensory attributes within the microalgal matrix.

### Bacterial strain selection based on bioactivity assays and dried weight yield

Building on the platform’s goal to enhance *C. vulgaris*’s suitability for food applications, a systematic screening pipeline was implemented to select bacterial strains capable of improving sensory and nutritional profiles through fermentation. Candidate strains were shortlisted based on food safety status (GRAS), literature evidence supporting their use in food fermentation, enzymatic potential relevant to flavour generation (**Figure 2A**), and preliminary sensory profiling. We referred to established safety frameworks to assess food-grade applicability, sourcing shortlisted candidate strains from the A*STAR NPL, a diverse repository for identifying strains with the potential for flavour enhancement and bioactive compound production. Alongside this, literature reviews were conducted to identify strains employed in food fermentations with known abilities to generate desirable flavour profiles and functional metabolites. Based on these criteria, a total of 22 strains spanning 13 microbial genera were selected. The most represented genera include *Bacillus* (18.2%), *Lacticaseibacillus* (13.6%), and *Lactiplantibacillus* (13.6%). The remaining strains belonged to *Cupriavidus, Microbacterium, Weizmannia*, and *Priestia*. The shortlisted panel also included other lactic acid bacteria (LAB) genera such as *Leuconostoc, Pediococcus, Ligilactobacillus, Liquorilactobacillus, Streptococcus*, and *Limosilactobacillus* (**Figure 2B**).

**Figure 2.**
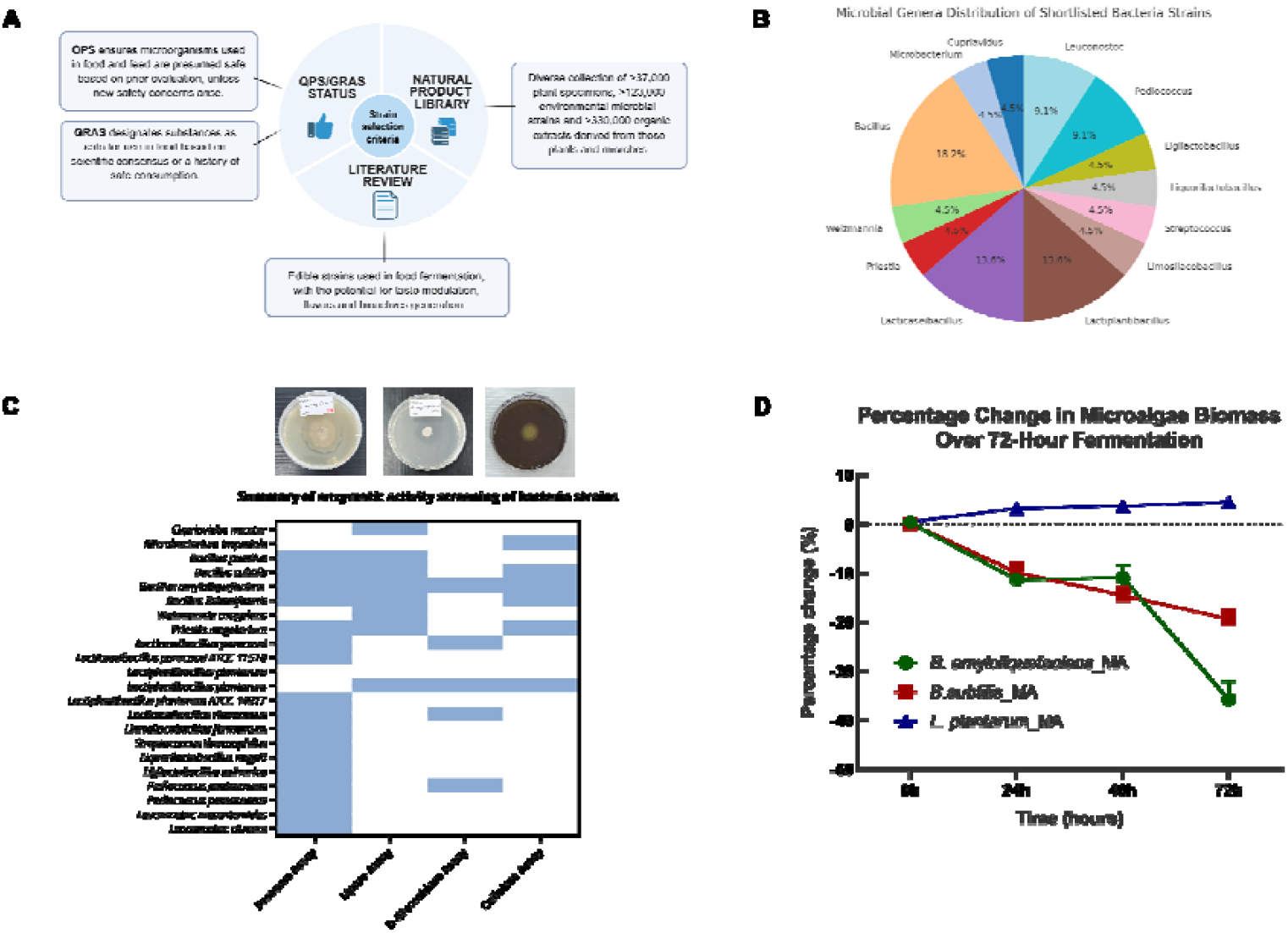
Strain selection and enzymatic characterization within the microbial biotransformation platform. **(A)** Candidate A*STAR NPL strains were selected based on food safety status (GRAS) and literature evidence supporting their use in food fermentation. **(B)** Distribution of shortlisted bacterial genera, with *Bacillus* (18.2%), *Lactiplantibacillus* (13.6%), and *Lacticaseibacillus* (13.6%) being the most dominant. **(C)** Enzymatic activities (protease, lipase, cellulase, β-glucosidase) were evaluated through qualitative and quantitative assays, with representative images and a heatmap summarising strain-specific enzymatic activity profiles. **(D)** Strain-dependent biomass changes in microalgae fermented with *B. amyloliquefaciens, B. subtilis*, and *L. plantarum* over a 72-hour time course relative to unfermented microalgae (control). *L. plantarum*-fermented samples retained or slightly increased biomass, while both *Bacillus* strains led to progressive biomass degradation, with *B. amyloliquefaciens* showing the most substantial loss by 72 hours.

The selected strains were subsequently tested in several enzymatic activity assays, including protease, lipase, β-glucosidase, and cellulase assays. Among the tested strains, *Bacillus amyloliquefaciens, Bacillus subtilis*, and *Lactiplantibacillus plantarum* demonstrated the highest levels of enzymatic activities (**Figure 2C, Table S1**). Even though *B. amyloliquefaciens* demonstrated superior enzymatic assay results across the board it also produced strong, undesirable odours that would significantly limit its inclusion rate and consumer appeal in a food matrix. Similarly, *B. subtilis* showed high enzymatic activity but was associated with biomass degradation and potential off flavours. On the other hand, while the selected *L. plantarum* strain displayed moderate enzymatic activity and did not show strong protease activity, it excelled in sensory attributes and well-documented health benefits crucial for functional food application. Furthermore, the absence of strong proteolytic activity is fairly advantageous, as excessive protein hydrolysis can lead to bitter peptide formation and off-flavors that compromise product acceptability. Moreover, microalgal proteins—such as those from *Chlorella* and *Spirulina*—are highly digestible and exhibit favourable amino acid profiles comparable to conventional protein sources ^12^. Crucially, this allows greater incorporation of the microalgal ingredient into food products, providing nutritional and functional health benefits without compromising sensory quality.

Given that the enzymatic activity profiles and preliminary sensory evaluation (in-house sniff tests) yielded divergent results regarding optimal strain performance, we decided to use dried weight yield analysis to evaluate the scalability and economic viability of the biotransformation platform by the shortlisted strains. We analysed the variations in dried weight during fermentation, a critical parameter often overlooked in microbial fermentation strategies, where substrate loss can undermine sustainability gains. The dry biomass yield of *Chlorella vulgaris* was monitored over 72 hours of fermentation with the three bacterial strains. Remarkably, *L. plantarum* fermentation resulted in a progressive biomass increase, gaining ∼5% by 72 hours, while *B. subtilis* and *B. amyloliquefaciens* caused substantial biomass losses of ∼15% and ∼40%, respectively (**Figure 2D**). This contrasting behaviour likely reflects fundamental differences in metabolic strategies: while *Bacillus* species catabolize algal biomass as the electron donor for respiration, *L. plantarum* demonstrates biomass-preserving metabolism for biosynthetic processes during aerobic fermentation, as previously reported for phylogenetically close *Lactobacilli* ^13^. The 45% differential in final biomass between *L. plantarum* (+5%) and *B. amyloliquefaciens* (−40%) represents a substantial economic advantage for industrial implementation, where feedstock costs typically dominate operational expenses. This dried weight enhancement by *L. plantarum* transforms the fermentation from a degradative process to an accumulative one, effectively amplifying the carbon-negative impact of the microalgae feedstock. Together, based on overall evaluation encompassing enzymatic activity profiles, preliminary sensory characteristics, and biomass preservation capacity, *L. plantarum* emerged as the optimal bacterial strain for *Chlorella vulgaris* biomass fermentation.

### Olfactory profile transformation of microalgae through bacterial fermentation

Subsequently, given the critical role of sensory attributes in consumer acceptance of microalgae-based foods, tiered olfactory profile tests and GC-MS analyses were designed to assess *L. plantarum*’s ability to transform *C. vulgaris*’s unpalatable sensory profile, building on its selection for enzymatic and sustainability benefits. We ran a time-course sniff test (n = 10 panellists) on *L. plantarum*-fermented microalgae samples (**Figures 3AB**). At baseline (0 hours), unfermented microalgae had a strong grassy odour, a known barrier to consumer acceptance. However, fermentation with *L. plantarum* notably shifted this profile. By 96 hours, the grassy odour had decreased markedly compared to unfermented controls.

**Figure 3.**
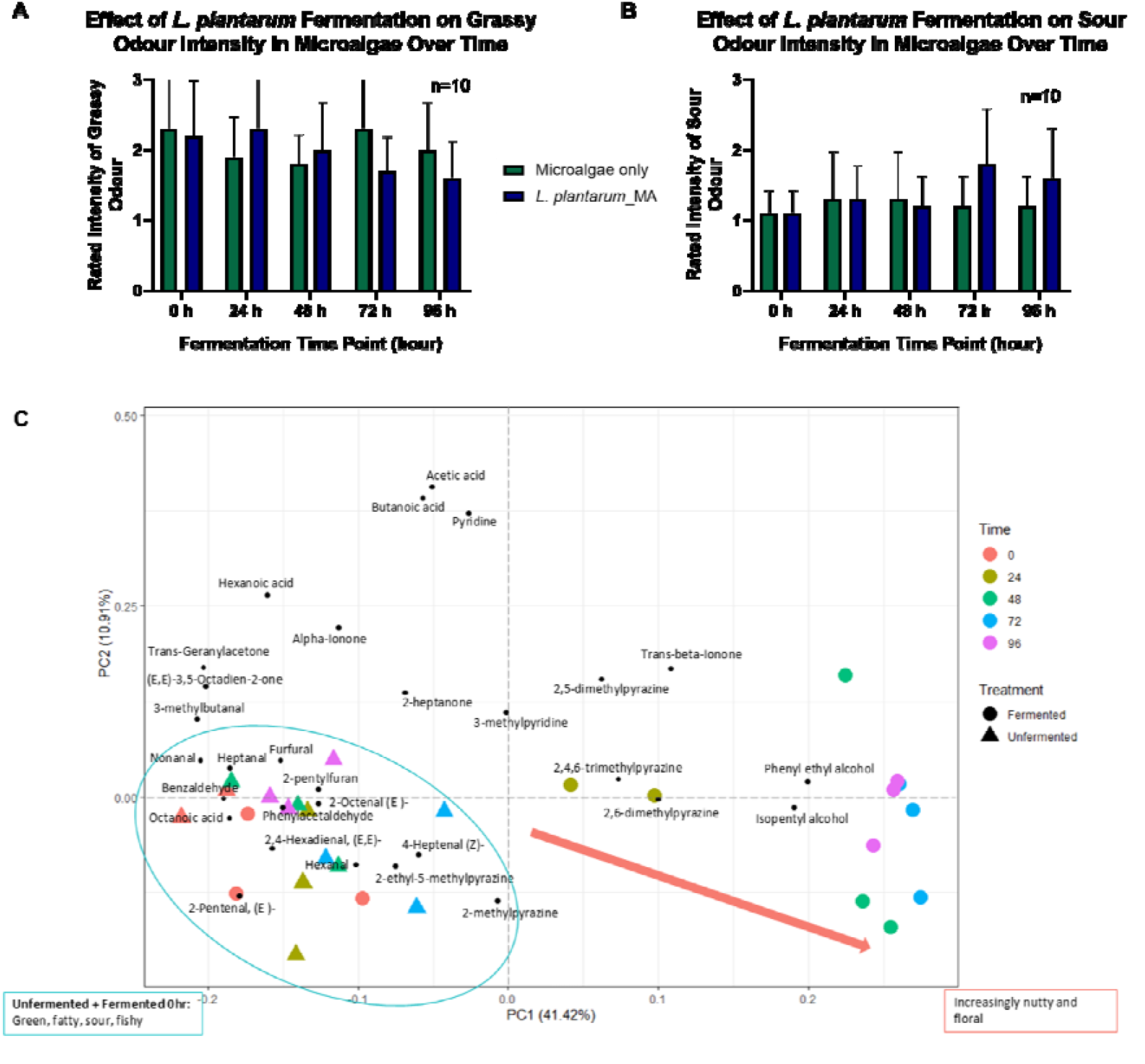
Olfactory and chemical profiling of microalgae fermented with *L. plantarum*. **(A-B)** Sensory intensities of **(A)** grassy and **(B)** sour odour attributes in unfermented microalgae (Microalgae only) and *L. plantarum*-fermented microalgae (*L. plantarum*_MA) over 96 hours. Olfactory evaluation was performed by an untrained in-house olfactory panel (n = 10), with intensities ranked on a structured 3-point scale (1 = weak, 2 = moderate, 3 = strong). Bars represent mean ± standard deviation. **(C)** Principal component analysis (PCA) analysis of volatile compounds identified by GC-MS in unfermented microalgae and *L. plantarum*-fermented microalgae, highlighting the temporal shift in aroma profile from green and fishy toward nutty and floral attributes as fermentation progresses.

To confirm this, we used a two-way repeated measures ANOVA—a statistical test that examines how two factors (here, fermentation treatment and time) influence an outcome (odour intensity), while accounting for repeated measurements from the same panellists. There was a significant interaction between treatment and time (P = 0.0188) for grassy odour between the fermented and unfermented samples (declining more in fermented ones). There were no significant main effects for time alone (P = 0.0845) or treatment alone (P = 0.6531), indicating neither factor independently drove the changes without the other. Panellists showed notable differences in their ratings (P<0.0001), reflecting natural variability in smell perception. Since the data violated the sphericity assumption (a requirement for ANOVA accuracy), we applied the Geisser-Greenhouse correction (ε = 0.8204) to adjust the results. Fermentation also increased sour notes over time. The same ANOVA test also revealed a significant interaction between treatment and time (P = 0.0287) for sour notes, showing that sourness evolved differently based on whether samples were fermented. No significant main effects appeared for time (P = 0.0607) or treatment (P = 0.3431) in isolation **(Tables S8-11**). Again, panellist variability was high (P<0.0001), and we used Geisser-Greenhouse correction for sphericity violation (ε = 0.5751). As negative controls, we assessed fishy and earthy odours, which showed no significant differences (P>0.05) (**Tables S12-15**) between fermented and unfermented samples (**Figure S3**), ruling out non-specific effects and strengthening the specificity of grassy and sour changes.

GC-MS analysis corroborated the sensory findings, providing molecular insights into the aroma transformation (**Figure 3C**). The volatile profile of microalgae fermented with *L. plantarum* revealed a significant decrease in polyunsaturated fatty acids (PUFAs)-derived aldehydes such as hexanal, 2-octenal, and 4-heptanol, which are typically associated with lipid oxidation and the undesirable “fishy” and “grassy” odours characteristic of microalgae ^14^. This reduction in off-flavours was accompanied by an increase in more desirable aroma compounds. Fermentation led to the accumulation of flavour-active compounds, including benzaldehyde, phenylethyl alcohol, pyrazines, and ionone derivatives. The increased production of pyrazines contributed to nutty and roasted notes ^15^, while phenylethyl compounds introduced floral aromas ^16^. Notably, pyrazines and ionone derivatives have also been identified as key volatiles in matcha, where they contribute to its roasted and floral– violet-like notes, respectively ^17^, highlighting their potential role in enhancing the sensory appeal of fermented microalgae. By 96 hours of fermentation, however, the aroma began to develop tangy sour notes of *L. plantarum*-fermented microalgae. PCA analysis, a technique that visualizes data clustering to show differences, of the volatile compounds showed a clear separation between control and fermented samples over time, indicating a dynamic shift in the volatile metabolome induced by bacterial activity. This separation became more pronounced with extended fermentation, highlighting the progressive nature of the sensory transformation. Together, these beneficial volatile profile modifications substantiate *L. plantarum’s* role in enhancing the sensory qualities of *Chlorella*, providing the foundation for subsequent taste and metabolite analyses.

### Taste modulation and metabolite profile evolution (E-tongue and LC-MS)

To further enhance *C. vulgaris*’s palatability and characterize its functional benefits, building on the sensory improvements achieved through *L. plantarum* fermentation, E-tongue and LC-MS analyses were conducted to evaluate taste modulation and metabolite changes. E-tongue analysis revealed a clear divergence between fermented samples and unfermented controls over the 96-hour fermentation period, particularly along principal component axes associated with umami and mild sourness (**Figure 4A**). The unfermented controls remained tightly clustered throughout, indicating a relatively stable taste profile and minimal variation in the absence of bacterial activity. In contrast, *L. plantarum*-fermented samples exhibited progressive shifts in the PCA space, particularly along PC1, reflecting dynamic bacterial-induced changes in taste-active metabolite composition. The distinct clustering of *L. plantarum*-fermented samples at later time points (48, 72, and 96 hours) suggests an active fermentation-driven metabolic transformation process that continuously alters the taste-active compound profile, while the unfermented controls remain compositionally stable. Overall, the E-tongue findings demonstrated that bacterial fermentation of microalgae leads to substantial shifts in taste properties. LC-MS analysis provided deeper insights into the specific metabolite transformations occurring during bacterial fermentation of microalgae. Fermentation with *L. plantarum* led to increased levels of umami-enhancing compounds, including glutamic acid and nucleotides such as adenosine monophosphate (AMP) and guanosine monophosphate (GMP) which are well-documented contributors to savoury taste perception (**Figure 4B**) ^18^.

**Figure 4.**
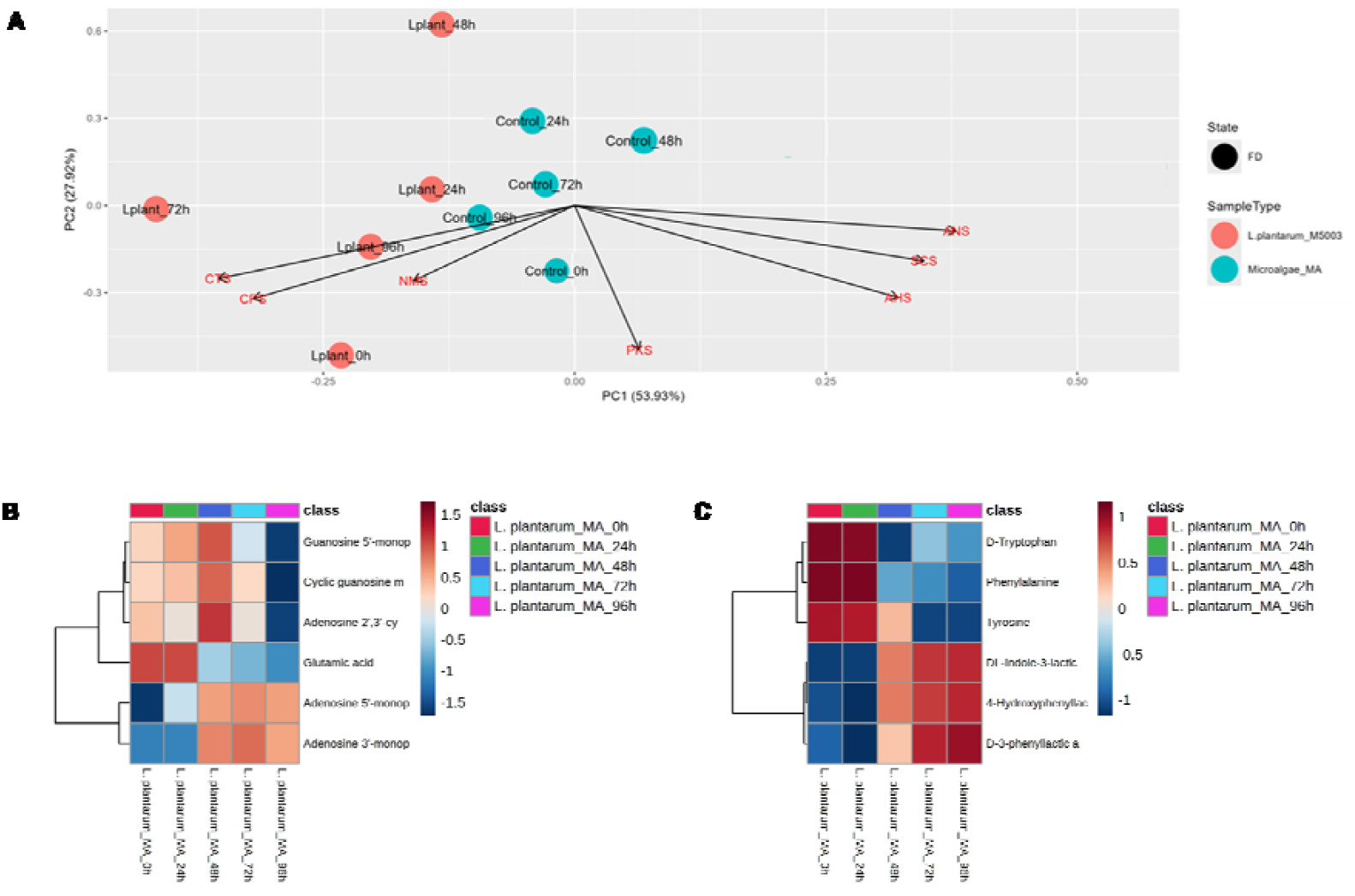
Multivariate and taste metabolite analysis of *L. plantarum*-fermented microalgae. **(A)** Principal component analysis (PCA) of E-tongue data showing temporal shifts in taste profiles of *L. plantarum*-fermented microalgae (*L. plantarum*_M5003) compared to unfermented microalgae (Microalgae_MA) over 96 hours. **(B)** LC-MS heatmap of selected taste-active metabolites, including glutamic acid, adenosine phosphates, and cyclic nucleotides, revealing dynamic accumulation patterns detected in *L. plantarum*-fermented microalgae *(L. plantarum*_MA) across fermentation. **(C)** LC-MS heatmap of aromatic amino acids (phenylalanine, L-tyrosine, D-tryptophan) and their derived metabolites across the same time course, indicating progressive conversion of precursors into downstream derivatives.

Additionally, the accumulation of three aromatic acids—indole lactic acid, phenyl lactic acid, and hydroxyphenyl lactic acid—was observed, with concurrent decreases in their amino acid precursors phenylalanine, L-tyrosine, and D-tryptophan (**Figure 4C**). These aromatic acids are known for their antioxidant and antimicrobial properties ^19^, suggesting that *L. plantarum* fermentation may confer ancillary functional benefits alongside sensory improvements. The metabolite profiles revealed by LC-MS analysis provided mechanistic explanations for the taste changes detected by the E-tongue. The increase in umami compounds in *L. plantarum*-fermented microalgae likely contributed to the umami-associated PCA shifts, while the production of organic acids possibly influenced the sourness dimension. Altogether, the taste and metabolite enhancements validate the ability of the probiotic-fermented microalgae to produce health-promoting, palatable food ingredients, supporting its sustainability evaluation.

## Conclusions

The biotransformation of microalgae *via* LAB fermentation offers a process-intensified, green alternative to conventional microalgae valorisation strategies. This study demonstrates probiotic fermentation by GRAS-certified *Lactiplantibacillus plantarum* as a sustainable, scalable biotransformation strategy for edible microalgae. Sensory and metabolomic profiling confirmed significant improvements in aroma, taste, and bioactive compound profiles. Compared to conventional microalgae valorisation methods such as solvent extraction, acid/base hydrolysis, and isoelectric precipitation—each associated with high E-factors (5– 30), chemical inputs, and significant energy demands—the probiotic fermentation platform developed in this study offers a substantial green advancement (**Table 1**). Operating under mild, aerobic conditions without solvents or pH-altering agents, the platform enables biomass preservation (+5%) while generating functional food ingredients directly from CO□-derived biomass. Crucially, the process is catalytic rather than stoichiometric: aromatic amino acids are converted into antioxidant lactates without degrading the bulk biomass. This fermentation avoids post-processing steps, maintains a carbon-negative profile, and valorises all biomass components. These attributes align with *Green Chemistry* principles 1 (waste prevention), 6 (energy efficiency), 9 (catalysis), and 10 (design for degradation), positioning this strategy as a model for sustainable food biomanufacturing.

**Table 1.** Green advances in microalgae processing using probiotics.

| **Green Chemistry Principle** | **Probiotic Fermentation** | **Chemical Processing** | **Enzymatic Processing** | **Green Advance & Literature Support** |
| --- | --- | --- | --- | --- |
| **1: Prevention** | Minimal organic waste | High chemical waste (*e.g.*, acidic/alkaline streams, salts) | Moderate enzyme waste | Reduced waste streams |
| **2: Atom Economy** | >95% theoretical yield | ~60–70% yields (with losses due to byproducts) | 80–85% yields | Higher atom incorporation |
| **3: Less Hazardous Synthesis** | Water and minerals only | Strong acids/bases (HCl, NaOH, KOH) | Mild enzyme solutions | Eliminates corrosive chemicals |
| **5: Safer Solvents** | Water-based system | Organic solvents required | Aqueous systems | Zero organic solvents |
| **6: Energy Efficiency** | 25–45°C | Often 100–370°C for thermal-chemical methods | 40–60°C | Mild temperature operation |
| **9: Catalysis** | Whole-cell biocatalyst | Chemical/thermal treatment | Isolated enzymes | Self-contained biocatalysis |

Data adapted from <sup>20-26</sup>.

The demonstrated biomass enhancement improves process economics by eliminating typical feedstock losses, while mild fermentation conditions (37 °C, 72 hours) are compatible with existing food fermentation infrastructure. Key scale-up considerations include maintaining consistent inoculum quality and optimizing aeration strategies to preserve the observed biomass-enhancing metabolism across different microalgae batches. In conclusion, through microalgae fermentation by *L. plantarum*, we achieved a reduction in off-flavour aldehydes, enhancement in umami compounds, and generation of bioactive metabolites—all without chemical additives or complicated downstream processing—thereby advancing microbial fermentation as both a sustainable bioprocessing platform and a practical route to circular food innovation.

## Glossary

(A*STAR NPL): Agency for Science, Technology, and Research’s Natural Product Library
(CO□): Carbon Dioxide
(E-tongue): Electronic Tongue
(GC-MS): Gas Chromatography-Mass Spectrometry
(GRAS): Generally Recognized As Safe
(LAB): Lactic Acid Bacteria
(LC-MS): Liquid Chromatography-Mass Spectrometry
(PCA): Principal Component Analysis
(PUFAs): Polyunsaturated Fatty Acids
(QPS): Qualified Presumption of Safety

## Author contributions

**Po-Hsiang Wang**: conceptualization, writing original draft, manuscript editing and supervision; **Zann Yi Qi Tan**: lab work, writing original draft and manuscript editing; **Choy Eng Nge**: lab work, manuscript review & editing; **Nurhidayah Basri**: lab work, manuscript review & editing; **Lina Xian Yu Lee**: manuscript writing, review & editing; **Aaron Thong**: manuscript review and editing; **Elaine Jinfeng Chin**: manuscript review & editing; **Sharon Crasta**: manuscript review and editing; **Geraldine Chan**: manuscript review & editing; **Mario Wibowo**: manuscript review and editing; **Yoganathan Kanagasundaram**: manuscript review & editing, supervision; **Siew Bee Ng**: manuscript review & editing, supervision. All authors have read and agreed to the published version of the manuscript.

## Supplementary Information

The supplementary information contains:

### Supplementary Materials and Methods

### Supplementary Figures

## Acknowledgments

This research is supported by the National Research Foundation, Prime Minister’s Office, Singapore, under its Campus for Research Excellence and Technological Enterprise (CREATE) programme [Proteins4Singapore] and Agency for Science, Technology and Research (A*STAR). We also thank Ryan Soh Han Wei for the use of his custom R script for the E-tongue analysis and Naedia Loh Xuan Yi for assisting in sample extraction and LCMS operations.

## References

1 C. L. Lumsden, J. Jägermeyr, L. Ziska and J. Fanzo, One Earth, 2024, 7, 1187–1201.

2 E. Daneshvar, R. J. Wicker, P.-L. Show and A. Bhatnagar, Chemical Engineering Journal, 2022, 427, 130884.

3 M. Chen, Y. Chen and Q. Zhang, Science of The Total Environment, 2024, 948, 174462.

4 F. Khavari, M. Saidijam, M. Taheri and F. Nouri, Molecular Biology Reports, 2021, 48, 4757–4765.

5 Y. Zhao, Q. Wang, D. Gu, F. Huang, J. Liu, L. Yu and X. Yu, Bioresource Technology, 2024, 393, 130093.

6 C. Chen, T. Tang, Q. Shi, Z. Zhou and J. Fan, Trends in Food Science & Technology, 2022, 126, 99–112.

7 C. Xue and I.-S. Ng, Bioresource Technology, 2022, 343, 126089.

8 F. Sandgruber, A. Gielsdorf, A. C. Baur, B. Schenz, S. M. Müller, T. Schwerdtle, G. I. Stangl, C. Griehl, S. Lorkowski and C. Dawczynski, Marine Drugs, 2021, 19, 310.

9 M. L. Olsen, K. Olsen and P. E. Jensen, Physiologia Plantarum, 2024, 176, e14337.

10 L. Machado, G. Carvalho and R. N. Pereira, Biomass, 2022, 2, 80–102.

11 F. L. Sirota, F. Goh, K.-N. Low, L.-K. Yang, S. C. Crasta, B. Eisenhaber, F. Eisenhaber, Y. Kanagasundaram and S. B. Ng, Journal of Genomics, 2018, 6, 63–73.

12 S. L. Ma, S. Sun, T. Z. Li, Y. J. Yan and Z. K. Wang, Critical Reviews in Food Science and Nutrition, 2024, 1–24.

13 M. J. A. Stevens, A. Wiersma, W. M. de Vos, O. P. Kuipers, E. J. Smid, D. Molenaar and M. Kleerebezem, Applied and Environmental Microbiology, 2008, 74, 4776–4778.

14 B. Coleman, C. Van Poucke, B. Dewitte, A. Ruttens, T. Moerdijk-Poortvliet, C. Latsos, K. De Reu, L. Blommaert, B. Duquenne, K. Timmermans, J. van Houcke, K. Muylaert and J. Robbens, Future Foods, 2022, 5, 100139.

15 F. Wang, H. Shen, T. Liu, X. Yang, Y. Yang and Y. Guo, Foods, 2021, 10, 273.

16 H. Issa-Issa, G. Guclu, L. Noguera-Artiaga, D. López-Lluch, R. Poveda, H. Kelebek, S. Selli and Á. A. Carbonell-Barrachina, Food Chemistry, 2020, 316, 126353.

17 Y. Luo, Y. Zhang, F. Qu, P. Wang, J. Gao, X. Zhang and J. Hu, Foods, 2022, 11, 2964.

18 J. Diepeveen, T. C. W. Moerdijk[Poortvliet and F. R. van der Leij, Journal of Food Science, 2022, 87, 1449–1465.

19 A. Bensid, N. El Abed, A. Houicher, J. M. Regenstein and F. Özogul, Critical Reviews in Food Science and Nutrition, 2020, 62, 2985–3001.

20 A. Kose and S. S. Oncel, Biotechnology Reports, 2015, 6, 137–143.

21 F. Almomani, H. Hosseinzadeh-Bandbafha, M. Aghbashlo, A. Omar, S.-W. Joo, Y. Vasseghian, H. Karimi-Maleh, S. Shiung Lam, M. Tabatabaei and S. Rezania, Chemical Engineering Journal, 2023, 455, 140588.

22 G. Karabulut, A. Purkiewicz and G. Goksen, Comprehensive Reviews in Food Science and Food Safety, 2024, 23, e13372.

23 L. Soto-Sierra, L. R. Wilken and C. K. Dixon, Bioresources and Bioprocessing, 2020, 7, 1–14.

24 O. Babich, S. Ivanova, P. Michaud, E. Budenkova, E. Kashirskikh, V. Anokhova and S. Sukhikh, Biotechnology Reports, 2025, 41, e00827.

25 P. S. Corrêa, W. G. Morais Júnior, A. A. Martins, N. S. Caetano and T. M. Mata, Processes, 2020, 9, 10.

26 S. Bleakley and M. Hayes, Foods, 2017, 6, 33.

