## Supporting information for "Sustainable biotransformation of microalgae *via* probiotic fermentation for enhanced functional, nutritional, and sensory properties"

#### Supporting Methods

#### Supporting Data

#### Supporting Tables

Table S1. Enzymatic activity profiles of 22 selected bacterial strains.

Table S2. Dry biomass yield (%w/v) and percentage change (%) of unfermented and fermented microalgae biomass over 72 hours.

Table S3. List of aromatic compounds detected in unfermented microalgae and *L. plantarum*-fermented microalgae over the course of fermentation.

Table S4. ASTREE E-tongue sensor summary with food applications.

Table S5. Metabolite profile of umami-related compounds detected during *L. plantarum*-mediated biotransformation of microalgae.

Table S6. Metabolite profile of amino acids and their bioconversion products during *L. plantarum*-mediated biotransformation of microalgae.

Table S7. Demographic and lifestyle information of the sniff test panelists (n = 10).

Table S8. Two-way ANOVA statistics for grassy odour intensity.

Table S9. Statistical summary of pairwise comparison for grassy aroma intensity between unfermented and in *L. plantarum*-fermented microalgae.

Table S10. Two-way ANOVA statistics for sour odour intensity.

Table S11. Statistical summary of pairwise comparison for sour odour intensity between unfermented and in *L. plantarum*-fermented microalgae.

Table S12. Two-way ANOVA statistics for fishy odour intensity.

Table S13. Statistical summary of pairwise comparison for fishy odour intensity between unfermented and in *L. plantarum*-fermented microalgae.

Table S14. Two-way ANOVA statistics for earthy odour intensity.

Table S15. Statistical summary of pairwise comparison for earthy odour intensity between unfermented and in *L. plantarum*-fermented microalgae.

#### Supporting Figures

Figure S1. LC-MS/MS fragmentation spectra at collision-induced dissociation (CID)10V of aromatic lactic acid derivatives in *L. plantarum*-fermented microalgae.

Figure S2. Extracted ion chromatograms (EICs) from LC-MS showing the bioconversion of precursor amino acids during *L. plantarum*-mediated fermentation of microalgae (MA) over 96 hours.

Figure S3. Olfactory evaluation of microalgae fermented with *L. plantarum.*

### Appendix

#### Supporting Methods

Unless specified otherwise, chemicals and reagents are purchased from Sigma-Aldrich (St. Louis, MO, USA).

##### ***Bacterial strains cultivation***

Non-lactic acid bacteria (non-LAB) strains—including *Bacillus* spp., *Cupriavidus necator*, *Microbacterium imperiale*, *Weizmannia coagulans*, and *Priestia megaterium*—were cultured in Luria-Bertani (LB) broth at 37 °C with agitation at 200 rpm until mid-logarithmic phase ($OD_{600}$ ≈ 0.4). Lactic acid bacteria (LAB) strains were cultivated in De Man, Rogosa, and Sharpe (MRS) broth under identical conditions (37 °C, 200 rpm) until $OD_{600}$ ≈ 0.4. *Streptococcus thermophilus* was cultured separately in M17 broth (Difco™, Becton, Dickinson and Company, Sparks, MD, USA) at 42 °C, 200 rpm till a similar growth phase.

##### ***Protease assay***

The proteolytic activity of the bacterial strains was qualitatively evaluated using the skim milk agar assay method. Skim milk agar was prepared with the following composition: 28 g/L skim milk powder, 5 g/L casein, 2.5 g/L yeast extract, 1 g/L dextrose, and 15 g/L agar in 1 L of distilled water, adjusted to a final pH of 7.0 ± 0.2 at 25 °C. 10 µL aliquot of the bacteria culture supernatant was aseptically spotted onto the surface of the agar and incubated at 37 °C. Proteolytic activity, as indicated by a zone of clearance around the inoculation site, was monitored across 96 hours.

***Lipase assay***

The lipolytic activity of the bacterial strains was qualitatively assessed using the tributyrin agar assay method. Tributyrin agar was prepared by dissolving 20 g of tributyrin agar powder in 1 L of distilled water, followed by adding 10 mL of glyceryl tributyrate upon cooling to 80 °C after autoclaving. 10 μL of the bacterial culture supernatant was spotted on the tributyrin agar and incubated at 37 °C. Lipolytic activity, as indicated by a zone of clearance around the inoculation site, was monitored across 96 hours.

***β-glucosidase assay***

β-glucosidase activity of the bacterial strains was assessed by quantifying the hydrolysis of *p*-nitrophenyl-β-D-glucopyranoside (pNPG). Bacterial culture supernatants were obtained by centrifugation at 4,000 × *g* for 20 minutes at 4 °C. For each sample, 0.5 mL of the culture supernatant was mixed with 2 mL of 1 mM pNPG solution and incubated at 45 °C for 30 minutes. The reaction was terminated by the addition of 2.5 mL of 1 M sodium carbonate. The release of *p*-nitrophenol was quantified by measuring absorbance at 400 nm using a spectrophotometer. β-glucosidase activity was calculated based on a standard calibration curve constructed using a commercial β-glucosidase enzyme, and all reactions were performed in triplicates.

***Cellulase assay***

Cellulolytic activity of the bacterial strains was qualitatively determined using the carboxymethyl cellulose (CMC) agar assay method. 0.2% CMC agar was prepared based on the following composition: CMC 2 g, sodium nitrate ($NaNO_3$) 20 mL, dipotassium hydrogen phosphate ($K_2HPO_4$) 1 g, potassium chloride (KCl) 1 g, magnesium sulfate ($MgSO_4$) 0.5 g, yeast extract 0.5 g, agar 15 g, 1 L distilled water. 10 μL of the culture supernatant was spotted on the 0.2% CMC agar and incubated at 37 °C. Cellulolytic activity, as indicated by a zone of clearance around the inoculation site, was monitored across 96 hours.

***Submerged fermentation of microalgae***

Submerged fermentation was conducted by inoculating *Chlorella vulgaris* with *L. plantarum* at an initial concentration of $10^6$ CFU/g. Each fermentation setup consisted of 5 g of sterile *C. vulgaris* biomass mixed with 50 mL of sterile distilled water in 250 mL Erlenmeyer flasks, with uninoculated *C. vulgaris* biomass as controls. Cultures were incubated at 37 °C with agitation at 200 rpm for up to 96 hours. Fermentates were sampled at five time points: 0 hours, 24 hours, 48 hours, 72 hours, and 96 hours. Harvested samples were immediately frozen at -80 °C and freeze-dried for 7 days. The dried fermentates were then ground into a fine powder using a sterile blender and stored at −20 °C until further analysis.

***Biomass yield and percentage change over time***

To determine the percentage dry yield of the biomass (% weight/volume), 0.5 mL of each sample of unfermented and fermented microalgae suspensions were pipetted into pre-weighed 5 mL tubes for each sample. The samples were frozen overnight in -80 °C freezer, followed by freeze-drying until fully dried. After drying, the tubes were weighed again to obtain the weight of the dry biomass. Data were analysed using GraphPad Prism version 8.0, and the percentage dry yield of the biomass (% weight/volume) and the percentage change in biomass (%) were calculated using the following formulae:

Dry Yield (%w/v) = (Weight of dry biomass (g) / Volume of wet sample (mL)) x 100

Percentage Change in Biomass (%) = [(Fermented biomass - Unfermented biomass) / (Unfermented biomass)] x 100

##### *In-house olfactory evaluations*

An in-house, untrained olfactory panel (n = 10) was recruited to perform time-course olfactory evaluations on microalgae fermentates inoculated with selected bacterial strains. Panelists assessed samples at various fermentation timepoints under blinded conditions, with all samples coded to eliminate bias. Aroma profiling focused on four key sensory attributes: grassy, sour, fishy, and earthy. Perceived odour intensity was rated using a structured 3-point scale, where 1 = *very weak*, 2 = *moderate*, and 3 = *very strong*. All evaluations were conducted anonymously. Panellists also completed a demographic and lifestyle questionnaire, which captured data on age, gender, ethnicity, dietary habits, smoking status, and self-rated smell sensitivity (on a 5-point scale, where 1 = *very poor* and 5 = *very strong*) (**Table S7**). Prior to participation, all individuals were briefed on the study objectives and procedures and provided written informed consent.

##### *Electronic tongue (E-tongue) analysis*

The taste and flavour evolution of the fermented microalgae samples was analysed using the ASTREE E-tongue system (Alpha MOS, Toulouse, France), a sensor-based technology designed to mimic human taste perception. The system employs seven potentiometric sensors (AHS, ANS, CPS, CTS, NMS, PKS, SCS) alongside an Ag/AgCl reference electrode, enabling the detection of key taste modalities such as bitterness, sweetness, umami, sourness, and saltiness, thereby generating a comprehensive taste fingerprint of each sample. E-tongue analysis was conducted following this protocol[1]. Briefly, 300 mg of freeze-dried fermentate was weighed into a 50 mL Falcon tube and reconstituted with 30 mL of sterile water. The mixture was vortexed and shaken on a rotary shaker for 30 minutes at room temperature, followed by centrifugation at 4000 × g for 30 minutes to obtain the supernatant. Between each sample run, the sensors and reference electrode were rinsed with water for 10 seconds to prevent cross-sample interference. Each sample was analysed for 120 seconds, with measurements repeated nine times to ensure system stability. The sensor signal at the 120th second was extracted from the last six measurements and averaged to generate the final raw data. Principal component analysis (PCA) was conducted using R software, and biplots were generated to visualize sample clustering and taste profile evolution during fermentation.

***Headspace Solid Phase Micro Extraction (HS-SPME) GC-MS Analysis***

Headspace solid-phase microextraction (HS-SPME) was conducted using a 23-gauge, 50/30 µm DVB/CAR/PDMS fibre (Supelco, USA). A volume of 500 µL of the wet sample was transferred into a 20 mL headspace vial and spiked with 100 µL of an internal standard solution (2,3-dimethoxytoluene at 10 ppm). The sample was incubated at 80 °C for 10 minutes, after which the SPME fibre was exposed to the headspace for 30 minutes while agitated at 250 rpm. Volatile compounds were then thermally desorbed from the fibre in the gas chromatograph

(GC) injector at 250 °C for 600 seconds. Subsequent GC-MS analysis was performed using an Agilent 7890B GC system coupled with a 5977B mass spectrometer and equipped with a multimode inlet (Agilent Technologies, USA). Separation was achieved on a polar J&W DB-Wax Ultra Inert capillary column (30 m × 0.25 mm I.D., 0.25 µm film thickness; Agilent Technologies). The injection was operated in split/splitless mode, with split ratios of 1:10. The oven temperature was initially set to 35 °C, then ramped at 100 °C/min to 60 °C, followed by 8 °C/min to 190 °C, and finally increased at 20 °C/min to 250 °C, where it was held for 6.5 minutes. Helium was used as the carrier gas at a constant flow of 1.0 mL/min. The mass spectrometer operated in electron ionization mode at 70 eV, with source and transfer line temperatures of 230 °C and 250 °C, respectively, and data was collected in full scan mode over the m/z range 35–450.

**Untargeted metabolite analysis by LC-MS**

Freeze-dried biomass (100 mg) was extracted with 20 mL of 50% aqueous MeOH. The mixture was vortexed briefly and sonicated for 10 min at room temperature. After sonication, the samples were centrifuged at 14,000 rpm for 5 min to separate the supernatant and solid biomass. The supernatant was transferred into LCMS glass vial for LCMS analysis (injection volume: 5 uL). Supernatant which exhibited turbidity were further filtered using 0.45 µm Claristep hydrophilic filter (Sartorius, Goettingen, Germany) prior to LCMS analysis.

HPLC-MS analyses were conducted using Agilent UHPLC 1290 Infinity coupled to Agilent 6540 accurate-mass quadrupole time-of-flight (QTOF) mass spectrometer and an ESI source. Gradient elution that starts from 98% water with 0.1% formic acid to 100% acetonitrile in 0.1% formic acid over 8.6 min along with an Acquity UPLC BEH C18 (2.1 × 50 mm, 1.7 µm) column at a flow rate of 0.5 mL/min was used. The operating parameters for QTOF were the same as previously reported [2].

The HRMS data were processed and analyzed using Agilent MassHunter Qualitative Analysis version 10.0 software; The untargeted metabolites analysis and annotation was performed using MZmine 3.9.0 [3]; Compounds annotation in MZmine was performed by matching experimental MS/MS spectra against library MS/MS spectra. The actual identity of indole lactic acid, phenyl lactic acid, and hydroxyphenyl lactic acid was validated by comparing MS/MS spectra and retention time with analytical standards. Metaboanalyst 6.0 was used for heatmap data visualization [4].

**Statistical analysis**

Olfactory evaluation data were analysed using GraphPad Prism version 8.0. For each aroma attribute, mean intensity scores were subjected to two-way repeated measures ANOVA, with treatment (unfermented vs. *Lactiplantibacillus plantarum*-fermented) and fermentation time-point (0, 24, 48, 72 and 96 hours) specified as within-subject factors. Panellist variation was included as a blocking factor to account for individual differences. The assumption of sphericity was not made, and Geisser-Greenhouse correction was applied where necessary. Statistical significance was determined at an alpha level of 0.05.

**Figures preparation**

BioRender (BioRender.com) was used to generate schematic figures for visualising experimental workflows and processes.

### Supporting Data

#### Supporting Tables

**Table S1. Enzymatic Activity Profiles of 22 Selected Bacterial Strains**

| Strains | Protease Activity | Lipase Activity | Cellulase Activity | β-Glucosidase Activity (mU/mL) |
|---|---|---|---|---|
| *Cupriavidus necator* | - | + | + | 0.05 |
| *Microbacterium imperiale* | - | - | + | 0 |
| *Bacillus pumilus* | +++ | ++ | - | 0 |
| *Bacillus subtilis* | ++ | ++ | +++ | 0.07 |
| *Bacillus amyloliquefaciens* | ++ | +++ | ++ | 0.73 |
| *Bacillus licheniformis* | + | ++ | ++ | 0.14 |
| *Weizmannia coagulans* | - | +++ | - | 0.14 |
| *Priestia megaterium* | +++ | ++ | + | 0 |
| *Lacticaseibacillus paracasei* | +++ | - | - | 0.40 |
| *Lacticaseibacillus paracasei ATCC 11578* | ++ | - | - | 0 |
| *Lactiplantibacillus plantarum* | - | - | - | 0 |
| *Lactiplantibacillus plantarum* | - | + | + | 0.48 |

| | | | | |
|---|---|---|---|---|
| *Lactiplantibacillus plantarum ATCC 14917* | +++ | - | - | 0 |
| *Lacticaseibacillus rhamnosus* | ++ | - | - | 0.54 |
| *Limosilacobacillus fermentum* | ++ | - | - | 0 |
| *Streptococcus thermophilus* | ++ | - | - | 0 |
| *Liquorilactobacillus nagelii* | ++ | - | - | 0 |
| *Ligilactobacillus salivarius* | ++ | - | - | 0 |
| *Pediococcus pentosaceus* | ++ | - | - | 0.22 |
| *Pediococcus pentosaceus* | ++ | - | - | 0 |
| *Leuconostoc mesenteroides* | ++ | - | - | 0 |
| *Leuconostoc citreum* | ++ | - | - | 0 |

Agar assays were used to assess protease, lipase, and cellulase activity and are scored qualitatively. Symbols represent relative activity levels: – (no activity), + (low), ++ (moderate), +++ (high). β-glucosidase activity was quantified from culture supernatants and expressed in milliunits per milliliter (mU/mL). Strains exhibiting **≥ 0.20mU/mL** of β-glucosidase activity are classified as positive (values highlighted in red).

**Table S2: Dry biomass yield (%w/v) and percentage change (%) of unfermented and fermented microalgae biomass over 72 hours**

| Sample | Dry Yield (% w/v) | | | Mean (n =3) | Standard deviation | Percentage change in fermented biomass (relative to unfermented controls) | | | Mean (n=3) | Standard deviation |
|---|---|---|---|---|---|---|---|---|---|---|
| | Replicate 1 | Replicate 2 | Replicate 3 | | | Replicate 1 | Replicate 2 | Replicate 3 | | |
| Microalgae_0h (unfermented) | 9.18 | 9.22 | 8.7 | 9.03 | 0.29 | - | - | - | - | - |
| Microalgae_24h (unfermented) | 8.92 | 9.16 | 9.24 | 9.11 | 0.17 | - | - | - | - | - |
| Microalgae_48h (unfermented) | 9.14 | 8.94 | 9.26 | 9.11 | 0.16 | - | - | - | - | - |
| Microalgae_72h (unfermented) | 9.4 | 9.1 | 9.08 | 9.19 | 0.18 | - | - | - | - | - |
| *B. amyloliquefaciens*_MA_0h | 9.2 | 9.26 | 8.72 | 9.06 | 0.30 | 0.22 | 0.43 | 0.23 | 0.29 | 0.12 |
| *B. amyloliquefaciens*_MA_24h | 7.94 | 8.16 | 8.16 | 8.09 | 0.13 | -10.99 | -10.92 | -11.69 | -11.20 | 0.43 |
| *B. amyloliquefaciens*_MA_48h | 8.38 | 7.72 | 8.26 | 8.12 | 0.35 | -8.32 | -13.65 | -10.80 | -10.92 | 2.67 |
| *B. amyloliquefaciens*_MA_72h | 6.22 | 5.46 | 6.04 | 5.91 | 0.40 | -33.83 | -40.00 | -33.48 | -35.77 | 3.67 |
| *B. subtilis*_MA_0h | 9.2 | 9.24 | 8.7 | 9.05 | 0.30 | 0.22 | 0.22 | 0.00 | 0.14 | 0.13 |
| *B. subtilis*_MA_24h | 8.18 | 8.32 | 8.14 | 8.21 | 0.09 | -8.30 | -9.17 | -11.90 | -9.79 | 1.88 |
| *B. subtilis*_MA_48h | 7.94 | 7.68 | 7.76 | 7.79 | 0.13 | -13.13 | -14.09 | -16.20 | -14.47 | 1.57 |
| *B. subtilis*_MA_72h | 7.7 | 7.42 | 7.14 | 7.42 | 0.28 | -18.09 | -18.46 | -21.37 | -19.30 | 1.80 |
| *L. plantarum*_MA_0h | 9.18 | 9.26 | 8.8 | 9.08 | 0.25 | 0.00 | 0.43 | 1.15 | 0.53 | 0.58 |
| *L. plantarum*_MA_24h | 9.2 | 9.46 | 8.98 | 9.21 | 0.24 | 3.14 | 3.28 | 3.22 | 3.21 | 0.07 |

| | | | | | | | | | | |
|---|---|---|---|---|---|---|---|---|---|---|
| *L. plantarum*_MA_48h | 9.46 | 9.28 | 9.6 | 9.45 | 0.16 | 3.50 | 3.80 | 3.90 | 3.73 | 0.21 |
| *L. plantarum*_MA_72h | 9.82 | 9.5 | 9.68 | 9.67 | 0.16 | 4.47 | 4.40 | 4.54 | 4.47 | 0.07 |

Dry biomass yield (% w/v) and percentage change in microalgae biomass (%) during 72-hour fermentation with selected microbial strains. *B. amyloliquefaciens*, *B, subtilis, L. plantarum* are labelled as "*B. amyloliquefaciens*_MA", "*B. subtilis*_MA" and "*L. plantarum*_MA" respectively; unfermented samples are labelled as Microalgae. Fermentation timepoints are expressed as 0h, 24h, 48h, and 72h, corresponding to 0, 24, 48, and 72 hours of fermentation, respectively. Dry biomass yield was quantified as dry weight (% w/v) and percentage change was calculated relative to unfermented controls at corresponding time points. Data represent mean ± standard deviation (n = 3 biological replicates). Notably, biomass decreased progressively during *Bacillus spp.* fermentations, with *B. amyloliquefaciens* showing the highest reduction (~36% at 72 h), while *L. plantarum* showed a modest increase in biomass over time (~+5%).

#### Table S3: List of volatiles detected in unfermented microalgae and *L. plantarum*-fermented microalgae throughout fermentation

| | | | | | | | | **Microalgae only (unfermented)** | | | | | ***L. plantarum*-fermented microalgae** | | | | |
|---|---|---|---|---|---|---|---|---|---|---|---|---|---|---|---|---|---|
| **No** | **Name** | **CAS No** | **Formula** | **Retention Time (min)** | **Retention Index (RI)** | **Compound class** | **Odour Description** | **0h** | **24h** | **48h** | **72h** | **96h** | **0h** | **24h** | **48h** | **72h** | **96h** |
| 1 | Acetic acid | 64-19-7 | C2H4O2 | 11.2 | 1462 | Acid | Sharp, sour, vinegar | 0.000373829 | 5.59499E-05 | 0.000102105 | 7.71635E-05 | 0.000117517 | 8.44834E-05 | 0.000230327 | 2.85498E-05 | 1.42339E-05 | 0 |
| 2 | Butanoic acid | 107-92-6 | C4H8O2 | 13.9 | 1640 | Acid | Sharp, acetic, cheese, dairy | 0.000224656 | 7.06529E-05 | 8.0395E-05 | 0.000100432 | 9.52012E-05 | 7.0737E-05 | 0.0002923 | 0 | 0 | 0 |
| 3 | Hexanoic acid | 142-62-1 | C6H12O2 | 17.0 | 1860 | Acid | Sour, fatty, sweaty, cheesy | 1.72991E-05 | 1.32659E-05 | 2.2313E-05 | 1.95154E-05 | 2.3045E-05 | 2.02186E-05 | 2.65836E-05 | 4.02418E-06 | 5.36998E-06 | 0 |
| 4 | Octanoic acid | 124-07-2 | C8H16O2 | 19.5 | 2077 | Acid | Fatty, waxy, rancid, cheesy | 1.73104E-05 | 4.65514E-05 | 3.47141E-05 | 2.71749E-05 | 5.11016E-05 | 4.19597E-05 | 2.47645E-05 | 0 | 0 | 0 |

| | | | | | | | | | | | | | | | | | |
|---|---|---|---|---|---|---|---|---|---|---|---|---|---|---|---|---|---|
| 5 | Propanoic acid | 79-09-4 | C3H6O2 | 12.6 | 1549 | Acid | Pungent, acidic, cheesy, vinegar | 0 | 0 | 0 | 1.44669E-05 | 1.71544E-05 | 0 | 1.33469E-05 | 0 | 0 | 0 |
| 6 | 1-Hexanol | 111-27-3 | C6H14O | 9.6 | 1358 | Alcohol | Green, fruity, sweet | 1.13061E-05 | 7.48597E-06 | 6.23502E-06 | 6.79656E-06 | 5.62038E-06 | 7.58196E-06 | 8.14862E-05 | 0 | 0 | 0 |
| 7 | 1-Octanol | 111-87-5 | C8H18O | 12.6 | 1548 | Alcohol | Mushroom, green, waxy | 0 | 0 | 0 | 0 | 0 | 0 | 0 | 0 | 0 | 4.31749E-06 |
| 8 | 1-Octen-3-ol | 3391-86-4 | C8H16O | 11.1 | 1453 | Alcohol | Mushroom | 0.000108602 | 7.89387E-05 | 0.00011246 | 0.000122192 | 0.000126945 | 0.000117102 | 0.000112986 | 5.94533E-05 | 4.57677E-05 | 4.48933E-05 |
| 9 | 1-Pentanol | 71-41-0 | C5H12O | 8.0 | 1256 | Alcohol | Fresh, bready, winey | 3.40815E-05 | 0.000108101 | 0.000111004 | 0.000106372 | 0.000110896 | 6.81737E-05 | 0.00016288 | 5.75653E-06 | 9.02769E-06 | 9.83897E-06 |
| 10 | 1-Penten-3-ol | 616-25-1 | C5H10O | 6.6 | 1163 | Alcohol | Pungent, green, horseradish, green vegetable, tropical, fruity | 0.000130451 | 0.000133269 | 8.80199E-05 | 0.000114489 | 0.000130628 | 9.5083E-05 | 7.84661E-05 | 0 | 0 | 0 |

| | | | | | | | | | | | | | | | | | |
|---|---|---|---|---|---|---|---|---|---|---|---|---|---|---|---|---|---|
| 11 | Isobutanol | 78-83-1 | C4H10O | 5.7 | 1098 | Alcohol | Ethereal, winey | 0 | 0 | 0 | 0 | 0 | 0 | 0 | 6.8378E-06 | 0 | 0 |
| 12 | Isopentyl alcohol | 123-51-3 | C5H12O | 7.4 | 1215 | Alcohol | Fusel, alcoholic, fruity, banana | 3.87013E-06 | 1.64211E-05 | 3.58926E-06 | 4.64628E-06 | 5.24433E-07 | 2.8019E-06 | 1.3368E-05 | 6.88474E-05 | 7.67948E-05 | 4.84116E-05 |
| 13 | Isophytol | 505-32-8 | C20H40O | 21.1 | 2290 | Alcohol | Floral, herbal, green | 0.000283568 | 0.000247227 | 0.00030138 | 0.000257763 | 0.000218515 | 0.000252264 | 0.000298784 | 0.000306844 | 0.00029257 | 0.000368416 |
| 14 | 2-Heptenal, (Z)- | 57266-86- | C7H12O | 9.2 | 1334 | Aldehyde | Green, fatty, sweet | 3.57133E-05 | 4.22325E-05 | 2.06006E-05 | 3.77043E-05 | 3.64459E-05 | 6.31658E-05 | 3.86303E-05 | 5.35539E-06 | 1.19544E-05 | 1.84186E-05 |
| 15 | 2-Hexenal, (E)- | 6728-26-3 | C6H10O | 7.6 | 1229 | Aldehyde | Green, banana, aldehydic, fresh, herbal | 4.15399E-05 | 2.95631E-05 | 4.40041E-05 | 3.84183E-05 | 4.39E-05 | 4.75577E-05 | 3.4967E-05 | 1.3438E-05 | 1.36717E-05 | 7.35864E-06 |
| 16 | 2-Nonenal, (E)- | 18829-56-6 | C9H16O | 12.6 | 1550 | Aldehyde | Cucumber, fatty, green | 1.3641E-05 | 1.93503E-05 | 6.58394E-06 | 1.08327E-05 | 1.23925E-05 | 7.66058E-06 | 4.31469E-06 | 5.83761E-06 | 2.55098E-06 | 0 |

| | | | | | | | | | | | | | | | |
|---|---|---|---|---|---|---|---|---|---|---|---|---|---|---|---|
| 17 | Heptanal | 111-71-7 | C7H14O | 7.0 | 1191 | Aldehyde | Fatty, green | 8.14218E-06 | 5.3004E-06 | 1.77838E-05 | 1.34867E-05 | 1.59664E-05 | 1.34607E-05 | 0 | 0 | 0 | 3.1134E-06 |
| 18 | Hexanal | 66-25-1 | C6H12O | 5.6 | 1085 | Aldehyde | Green, grassy | 2.95069E-05 | 4.9805E-05 | 7.3148E-05 | 3.17118E-06 | 4.81524E-05 | 8.81279E-05 | 2.97348E-06 | 2.18273E-05 | 2.39616E-05 | 1.65279E-05 |
| 19 | Nonanal | 124-19-6 | C9H18O | 10.3 | 1400 | Aldehyde | Green, cucumber | 3.67461E-05 | 3.55442E-05 | 3.17581E-05 | 3.06157E-05 | 2.04399E-05 | 2.30536E-05 | 9.41831E-06 | 8.65151E-06 | 1.49292E-05 | 1.39231E-05 |
| 20 | 2-Octenal, (E)- | 2548-87-0 | C8H14O | 10.9 | 1441 | Aldehyde | Fresh, cucumber, fatty, green, citrus | 3.0478E-05 | 3.47488E-05 | 2.22665E-05 | 3.24509E-05 | 3.86952E-05 | 2.44124E-05 | 0 | 9.47891E-06 | 1.05135E-05 | 9.66193E-06 |
| 21 | 2-Pentenal, (E)- | 1576-87-0 | C5H8O | 6.3 | 1138 | Aldehyde | Pungent, green, fruity, apple orange, tomato | 1.51164E-05 | 4.7341E-05 | 4.86063E-05 | 4.22268E-05 | 4.86639E-05 | 4.97087E-05 | 2.66813E-05 | 6.98378E-06 | 9.204E-06 | 2.39077E-05 |
| 22 | 3-methylbutanal | 590-86-3 | C5H10O | 3.9 | 921 | Aldehyde | Ethereal, aldehydic, chocolate, peach, fatty | 2.55363E-05 | 1.69893E-05 | 1.53889E-05 | 9.52913E-06 | 1.05045E-05 | 2.14092E-05 | 5.1961E-06 | 0 | 0 | 0 |

| | | | | | | | | | | | | | | | | |
|---|---|---|---|---|---|---|---|---|---|---|---|---|---|---|---|---|
| 23 | 4-Heptenal, (Z)- | 6728-31-0 | C7H12O | 7.9 | 1247 | Aldehyde | Green, oily, fatty, dairy | 3.67575E-06 | 5.52971E-06 | 4.70133E-06 | 5.25069E-06 | 5.82723E-06 | 5.47214E-06 | 3.67406E-06 | 4.42002E-06 | 0 | 2.93683E-06 |
| 24 | beta-Cyclocitral | 432-25-7 | C10H16O | 14.0 | 1643 | Aldehyde | Tropical, saffron, herbal, clean, rose, sweet, tobacco, green, fruity | 9.11098E-05 | 7.51192E-05 | 3.68818E-05 | 9.54359E-05 | 6.1081E-05 | 3.99324E-05 | 0.000107971 | 9.15599E-05 | 2.87444E-05 | 4.79827E-05 |
| 25 | cis-2-Butenal | 590-18-1 | C4H6O | 5.1 | 1047 | Aldehyde | Pungent, suffocating, irritating | 4.05156E-05 | 4.18626E-05 | 4.11305E-05 | 4.13284E-05 | 4.00872E-05 | 3.88505E-05 | 2.45241E-05 | 4.86375E-06 | 4.81032E-06 | 1.94214E-06 |
| 26 | Dodecanal | 112-54-9 | C12H24O | 15.3 | 1732 | Aldehyde | Soapy, waxy, aldehydic, citrus, green, floral | 0.000104569 | 4.85737E-06 | 1.72808E-05 | 0 | 1.15631E-05 | 5.45698E-06 | 4.29987E-06 | 4.93151E-06 | 0 | 2.65449E-06 |
| 27 | Octanal | 124-13-0 | C8H16O | 8.6 | 1296 | Aldehyde | Orange peel, green | 0 | 3.71881E-06 | 3.18694E-06 | 6.57751E-06 | 3.00568E-06 | 3.68664E-06 | 0 | 0 | 0 | 0 |
| 28 | Pentanal | 110-62-3 | C5H10O | 4.5 | 984 | Aldehyde | Fermented, bready, cocoa | 0.000164836 | 0.000127786 | 0.000128118 | 0.000111268 | 0.000125985 | 0.000137583 | 3.51212E-05 | 0 | 1.45685E-05 | 3.46825E-05 |

| | | | | | | | | | | | | | | | |
|---|---|---|---|---|---|---|---|---|---|---|---|---|---|---|---|
| 29 | Undecanal | 112-44-7 | C11H22O | 13.6 | 1614 | Aldehyde | Waxy, soapy, floral, aldehydic, citrus, green, fatty, laundered cloth | 0 | 3.1398E-06 | 2.86654E-06 | 0 | 0 | 0 | 0 | 0 | 0 | 0 |
| 30 | 4-ethylbenzaldehyde | 4748-78-1 | C9H10O | 15.2 | 1729 | Aromatic | Almond, bitter, sweet, anise cherry | 1.90247E-05 | 6.79073E-05 | 8.25158E-05 | 8.93263E-05 | 0.000107134 | 3.87547E-05 | 0 | 0 | 0 | 0 |
| 31 | Benzaldehyde | 100-52-7 | C7H6O | 12.4 | 1540 | Aromatic | Almond, cherry, sharp, sweet | 0.000686077 | 0.000670456 | 0.000738445 | 0.000639122 | 0.000723182 | 0.000667591 | 0.000174753 | 0.000287285 | 0.000264358 | 0.000245761 |
| 32 | Benzaldehyde, 2,4-dimethyl- | 613-45-6 | C9H10O | 15.5 | 1751 | Aromatic | Naphthyl, cherry, almond, spicy, vanilla | 0.0001 4304 | 0.00020615 | 0.000191227 | 0.00017273 | 0.000183541 | 0.000185366 | 6.22234E-05 | 3.82608E-05 | 1.39722E-05 | 3.62033E-05 |
| 33 | Benzaldehyde, 3-methyl- | 620-23-5 | C8H8O | 13.9 | 1641 | Aromatic | Sweet, fruity, cherry, almond | 3.63298E-05 | 0 | 0 | 0 | 0 | 0 | 0 | 0 | 0 | 0 |
| 34 | Benzene, 1,3-bis(1,1-dimethylethyl)- | 1014-60-4 | C14H22 | 10.8 | 1433 | Aromatic | - | 4.89165E-05 | 5.79204E-05 | 6.22572E-05 | 5.59074E-05 | 4.92305E-05 | 7.00738E-05 | 0 | 3.06168E-05 | 4.80208E-05 | 3.11789E-05 |

| | | | | | | | | | | | | | | | | | |
|---|---|---|---|---|---|---|---|---|---|---|---|---|---|---|---|---|---|
| 35 | Benzyl alcohol | 100-51-6 | C7H8O | 17.5 | 1892 | Aromatic | Floral, rose, phenolic, balsamic | 0 | 0 | 0 | 0 | 0 | 0 | 3.24128E-05 | 2.45013E-05 | 0 | 0 |
| 36 | Naphthalene, 1,2-dihydro-1,1,6-trimethyl- | 447-53-0 | C13H16 | 15.8 | 1768 | Aromatic | Licorice | 0.000171255 | 8.07668E-05 | 0.000194913 | 0.000154746 | 0.000165185 | 8.17133E-05 | 0.000179282 | 0.000128624 | 0.000209427 | 0.000213828 |
| 37 | Phenylacetaldehyde | 122-78-1 | C8H8O | 14.2 | 1659 | Aromatic | Rose, sweet, floral | 0 | 1.34409E-05 | 2.36096E-05 | 0 | 7.33735E-06 | 2.18227E-05 | 0 | 0 | 0 | 0 |
| 38 | Phenylethyl alcohol | 60-12-8 | C8H10O | 17.9 | 1933 | Aromatic | Floral, rose, sweet | 0.000291961 | 0.00025571 | 0.000275207 | 0.000254718 | 0.000274726 | 0.000177831 | 0.000301547 | 0.001545977 | 0.00238809 | 0.002478273 |
| 39 | p-Xylene | 106-42-3 | C8H10 | 6.4 | 1147 | Aromatic | Sweet | 1.11668E-05 | 0 | 0 | 0 | 0 | 8.23851E-06 | 0 | 0 | 0 | 0 |
| 40 | 2,4-Decadienal, (E,E)- | 25152-84-5 | C10H16O | 16.6 | 1827 | Enal | Satty, chicken | 0 | 0 | 0 | 0 | 1.28238E-05 | 0 | 0 | 0 | 0 | 0 |

| | | | | | | | | | | | | | | | |
|---|---|---|---|---|---|---|---|---|---|---|---|---|---|---|---|
| 41 | 2,4-Heptadienal, (E,E)- | 4313-03-5 | C7H10O | 11.9 | 1507 | Enal | Fatty, green, oily, aldehydic, vegetable, cinnamon | 0.000576431 | 0.000709327 | 0.000778931 | 0.00072895 | 0.000928734 | 0.000724294 | 0.000373158 | 0 | 5.45677E-05 | 0.000106013 |
| 42 | 2,4-Hexadienal, (E,E)- | 142-83-6 | C6H8O | 10.5 | 1415 | Enal | Sweet, green, spicy, floral, citrus | 2.61753E-05 | 4.4413E-05 | 3.19583E-05 | 1.63111E-05 | 1.95492E-05 | 5.34876E-05 | 0 | 4.86428E-06 | 3.27238E-06 | 8.94491E-06 |
| 43 | 2,4-Nonadienal, (E,E)- | 5910-87-2 | C9H14O | 15.0 | 1717 | Enal | Fatty, melon, cucumber, chicken fat | 1.71847E-05 | 7.44766E-05 | 7.43974E-05 | 5.47693E-05 | 9.9412E-05 | 8.66124E-05 | 1.7384E-05 | 0 | 0 | 9.03327E-06 |
| 44 | 2,6-Nonadienal, (E,Z)- | 557-48-2 | C9H14O | 13.3 | 1599 | Enal | Green, fatty, dry, cucumber, violet leaf | 4.21532E-05 | 2.43174E-05 | 2.53043E-05 | 3.00407E-05 | 2.21378E-05 | 1.305E-05 | 2.79858E-05 | 1.36566E-05 | 1.38285E-05 | 4.88694E-06 |
| 45 | Methyl palmitate | 112-39-0 | C17H34O2 | 20.7 | 2228 | Ester | Fatty, waxy | 6.4327E-05 | 1.85663E-05 | 3.67519E-05 | 5.34491E-05 | 2.14683E-05 | 3.95351E-05 | 2.32069E-05 | 4.1457E-05 | 4.10557E-05 | 1.84766E-05 |
| 46 | Methyl salicylate | 119-36-8 | C8H8O3 | 16.2 | 1799 | Ester | Wintergreen, minty | 0.000748143 | 0.000647308 | 0.000582888 | 0.000574573 | 0.000528463 | 0.000589075 | 0.000615054 | 0.000173008 | 0.000535453 | 0.000540567 |

| | | | | | | | | | | | | | | | | | |
|---|---|---|---|---|---|---|---|---|---|---|---|---|---|---|---|---|---|
| 47 | 2-Acetyl-5-methylfuran | 1193-79-9 | C7H8O2 | 13.4 | 1607 | Furan | Sweet, musty, nutty, caramellic | 0 | 0 | 0 | 0 | 0 | 0 | 0 | 0 | 8.27736E-05 | 0 |
| 48 | 2-ethylfuran | 3208-16-0 | C6H8O | 4.2 | 955 | Furan | Chemical, beany, ethereal, cocoa, bready, malty, coffee, nutty | 2.98867E-05 | 2.66821E-05 | 2.75321E-05 | 2.68418E-05 | 3.24093E-05 | 3.1151E-05 | 2.30977E-05 | 2.12433E-05 | 1.55444E-05 | 2.42648E-05 |
| 49 | 2-Pentylfuran | 3777-69-3 | C9H14O | 7.6 | 1234 | Furan | Fruity, beany, vegetable | 0.000262495 | 0.000170152 | 0.000153915 | 0.000109902 | 8.60189E-05 | 0.000304494 | 8.22013E-05 | 7.5605E-05 | 6.40852E-05 | 0.000113595 |
| 50 | 3-Furanmethanol | 4412-91-3 | C5H6O2 | 14.3 | 1669 | Furan | Sweet, nutty, herbal | 0 | 0 | 0 | 0 | 0 | 0 | 1.66561E-05 | 0 | 0 | 0 |
| 51 | 5-methyl furfural | 620-02-0 | C6H6O2 | 12.9 | 1568 | Furan | Sweet, caramel, maple | 1.64153E-05 | 1.40762E-05 | 1.45381E-05 | 1.31659E-05 | 1.43611E-05 | 1.52432E-05 | 0 | 0 | 0 | 0 |
| 52 | Furan, 2-methyl- | 534-22-5 | C5H6O2 | 3.6 | 874 | Furan | Chocolate, ethereal, acetone | 3.37885E-06 | 1.99509E-06 | 1.92152E-06 | 1.74903E-06 | 0 | 3.19082E-06 | 5.8813E-07 | 1.14447E-06 | 0 | 1.37179E-06 |

| | | | | | | | | | | | | | | | |
|---|---|---|---|---|---|---|---|---|---|---|---|---|---|---|---|
| 53 | Furfural | 98-01-1 | C5H4O2 | 11.4 | 1473 | Furan | Sweet, woody, almond, bread, baked | 0.000192257 | 0.000176687 | 0.000234249 | 0.000142119 | 9.04195E-05 | 0.000169261 | 2.00106E-05 | 2.41157E-06 | 0 | 0 |
| 54 | trans-2-(2-Pentenyl)furan | 70424-14-5 | C9H12O | 8.8 | 1306 | Furan | Bean, grassy | 5.52195E-06 | 1.06584E-05 | 1.05849E-05 | 9.12875E-06 | 1.08564E-05 | 9.22086E-06 | 0 | 1.83451E-06 | 3.56662E-07 | 0 |
| 55 | 1,3,5,7-Cyclooctatetraene | 629-20-9 | C8H8 | 8.1 | 1265 | Hydrocarbon | - | 0 | 0 | 0 | 0 | 0 | 0 | 0 | 0 | 0 | 1.98204E-06 |
| 56 | 3-Heptadecene, (Z)- | 68155-00-0 | C9H10O | 15.1 | 1723 | Hydrocarbon | - | 0.000283823 | 0.000362702 | 0.00058726 | 0.000529519 | 0.00050902 | 0.000171216 | 0.000183636 | 0.000484612 | 0.000403239 | 0.000484425 |
| 57 | Heptadecane | 629-78-7 | C17H36 | 14.8 | 1698 | Hydrocarbon | Waxy | 0.000700864 | 0.000731208 | 0.001215075 | 0.001006742 | 0.001170633 | 0 | 0.000442546 | 0.00047186 | 0.000493414 | 0.000950684 |
| 58 | Pentadecane | 629-62-9 | C15H32 | 11.8 | 1498 | Hydrocarbon | - | 4.04429E-05 | 9.63707E-05 | 0.000130275 | 9.38504E-05 | 0.000123106 | 6.7386E-05 | 6.13493E-05 | 4.87533E-05 | 5.03987E-05 | 5.40487E-05 |

| | | | | | | | | | | | | | | | | | |
|---|---|---|---|---|---|---|---|---|---|---|---|---|---|---|---|---|---|
| 59 | Undecane | 1120-21-4 | C10H22 | 4.6 | 999 | Hydrocarbon | - | 0 | 4.80151E-06 | 5.10924E-06 | 4.5991E-06 | 0 | 3.23377E-06 | 0 | 2.66008E-06 | 0 | 6.42428E-06 |
| 60 | (E,E)-3,5-Octadien-2-one | 30086-02-3 | C8H12O | 13.1 | 1584 | Ketone | Fruity, green, grassy | 0.001027837 | 0.000853237 | 0.00081154 | 0.00071048 | 0.000752466 | 0.000970566 | 0.000856894 | 0.000211837 | 0.000106335 | 7.81424E-05 |
| 61 | 1-Octen-3-one | 4312-99-6 | C8H14O | 8.8 | 1309 | Ketone | Herbal, mushroom, earthy, must,y dirty | 1.3969E-05 | 1.60312E-05 | 1.79359E-05 | 0 | 1.32361E-05 | 1.37119E-05 | 1.8517E-05 | 1.60293E-05 | 5.74918E-06 | 1.69357E-05 |
| 62 | 1-Penten-3-one | 1629-58-9 | C5H8O | 4.9 | 1026 | Ketone | Spicy, pungent, peppery, mustard, garlic, onion | 0 | 0 | 1.80323E-05 | 0 | 0 | 0 | 1.4325E-05 | 0 | 0 | 0 |
| 63 | 1-Propanone, 1-(2-furanyl)- | 3194-15-8 | C7H8O2 | 13.2 | 1590 | Ketone | Fruity | 0 | 0 | 0 | 0 | 0 | 0 | 0 | 0 | 3.11861E-06 | 4.629E-06 |
| 64 | 2,3-Butanedione | 431-03-8 | C4H6O2 | 4.4 | 976 | Ketone | Buttery, sweet, creamy, pungent, caramellic | 0 | 0 | 0 | 1.55107E-05 | 0 | 0 | 0 | 2.16852E-05 | 0 | 0 |

| | | | | | | | | | | | | | | | | | |
|---|---|---|---|---|---|---|---|---|---|---|---|---|---|---|---|---|---|
| 65 | 2,3-Pentanedione | 600-14-6 | C5H8O2 | 5.2 | 1056 | Ketone | Sweet, buttery, creamy | 7.004E-06 | 2.19717E-05 | 2.35508E-05 | 2.08714E-05 | 2.55922E-05 | 2.23478E-05 | 1.63437E-05 | 1.83899E-05 | 1.67855E-05 | 5.24942E-06 |
| 66 | 2-Butanone | 78-93-3 | C4H8O | 3.8 | 908 | Ketone | Acetone, ethereal, fruity, camphoraceous | 0.00024514 | 0.000221226 | 0.000348183 | 0.000155413 | 0.000153143 | 0.000223449 | 0.000126828 | 0.000274341 | 0.000202228 | 0.000144416 |
| 67 | 2-Heptanone | 110-43-0 | C7H14O | 7.0 | 1187 | Ketone | Cheesy, fruity, ketogenic, green, banana, creamy | 3.44669E-05 | 1.6017E-05 | 1.51541E-05 | 0 | 0 | 0 | 0 | 0 | 0 | 0 |
| 68 | 2-Nonanone | 821-55-6 | C9H18O | 10.2 | 1397 | Ketone | Fruity, sweet, waxy, soapy, cheesy, green, herbal, coconut | 1.36669E-05 | 2.43512E-05 | 3.91895E-05 | 0 | 0 | 4.48629E-05 | 1.56326E-05 | 0 | 0 | 0 |
| 69 | 2-Pentadecanone, 6,10,14-trimethyl- | 502-69-2 | C18H36O | 20.0 | 2137 | Ketone | Oily, herbal, jasmine, celery, woody | 0.000321032 | 0.000296771 | 0.000322788 | 0.000317161 | 0.000394462 | 0.000299194 | 0.000207134 | 0.000187917 | 0.000234206 | 0.000212284 |
| 70 | 2-Tridecanone | 593-08-8 | C13H26O | 16.5 | 1820 | Ketone | Fatty, waxy, dairy, milky | 2.19476E-05 | 2.93369E-05 | 1.91158E-05 | 2.79608E-05 | 3.16983E-05 | 2.98741E-05 | 2.02939E-05 | 0 | 0 | 0 |

| | | | | | | | | | | | | | | | | | |
|---|---|---|---|---|---|---|---|---|---|---|---|---|---|---|---|---|---|
| 71 | 3,5-Octadien-2-one | 38284-27-4 | C8H12O | 12.3 | 1532 | Ketone | Fruity, fatty, mushroom | 0.000141213 | 0.00015384 | 0.000166226 | 9.56053E-05 | 0.00012918 | 0.000108819 | 9.72888E-05 | 0 | 0 | 0 |
| 72 | 3-Nonen-2-one | 14309-57-0 | C9H16O | 12.2 | 1524 | Ketone | Fruity, berry, fatty, oily | 0 | 1.54487E-05 | 1.45496E-05 | 2.19196E-05 | 2.7222E-05 | 2.13097E-05 | 1.19724E-05 | 5.08251E-06 | 0 | 0 |
| 73 | 3-Octanone | 106-68-3 | C8H16O | 8.1 | 1260 | Ketone | Lavender, mushroom, sweet | 1.00294E-05 | 7.74798E-06 | 3.31775E-06 | 0 | 0 | 9.17242E-06 | 2.68099E-05 | 0 | 0 | 1.46366E-06 |
| 74 | 3-Octanone, 2-methyl- | 923-28-4 | C9H18O | 9.0 | 1323 | Ketone | - | 0.00010475 | 7.37462E-05 | 0.000104964 | 0 | 3.45306E-05 | 0.000123455 | 1.74983E-05 | 0 | 0 | 0 |
| 75 | 5-Hepten-2-one, 6-methyl- | 110-93-0 | C8H14O | 9.4 | 1345 | Ketone | Citrus, green, musty, lemongrass, apple | 2.80764E-05 | 2.42381E-05 | 2.37667E-05 | 2.75072E-05 | 3.0203E-05 | 2.55302E-05 | 1.94399E-05 | 0 | 0 | 0 |
| 76 | 6-Methyl-3,5-heptadiene-2-one | 1604-28-0 | C8H12O | 13.5 | 1608 | Ketone | Spicy, cinnamon, coconut, spicy, woody, sweet, weedy | 0.00018724 | 0.000245117 | 0.000243097 | 0.000150534 | 0.000235556 | 0.000263096 | 0.000179241 | 0 | 0 | 0 |

| | | | | | | | | | | | | | | | | | |
|---|---|---|---|---|---|---|---|---|---|---|---|---|---|---|---|---|---|
| 77 | Acetoin | 513-86-0 | C4H8O2 | 8.7 | 1299 | Ketone | Buttery, creamy, dairy, milky, fatty | 0 | 0 | 0 | 0 | 1.11621E-05 | 0 | 0 | 0 | 0 | 0 |
| 78 | Acetone | 67-64-1 | C3H6O | 3.2 | 818 | Ketone | Solvent, ethereal, apple, pear | 0 | 0 | 0 | 0 | 1.89788E-05 | 0 | 1.27531E-05 | 1.87863E-05 | 9.59586E-05 | 9.93838E-05 |
| 79 | Acetophenone | 98-86-2 | C8H8O | 14.4 | 1670 | Ketone | Sweet, hawthorn | 1.47569E-05 | 0 | 0 | 0 | 0 | 0 | 1.45063E-05 | 0 | 0 | 0 |
| 80 | Cyclohexanone, 2,2,6-trimethyl- | 2408-37-9 | C9H16O | 9.2 | 1335 | Ketone | Thujonic, pungent, labdanum, honey | 2.48519E-05 | 5.27998E-05 | 4.69297E-05 | 2.01988E-05 | 3.13161E-05 | 4.28309E-05 | 2.55285E-05 | 2.17541E-05 | 0 | 5.135E-06 |
| 81 | Isophorone | 78-59-1 | C9H14O | 13.6 | 1621 | Ketone | Cooling, woody, sweet, green, camphoraceous, fruity, musty | 8.98165E-05 | 0.000126727 | 4.45773E-05 | 0 | 4.33334E-05 | 3.9674E-05 | 8.85449E-05 | 4.15323E-05 | 7.95499E-05 | 0.000117045 |
| 82 | m-Methylacetophenone | 585-74-0 | C9H10O | 16.2 | 1795 | Ketone | Hawthorn, sweet, mimosa, coumarinic, cherry, acacia | 0.000203678 | 0.000213941 | 0.000214632 | 0.00019917 | 0.000202363 | 0.000236624 | 0.000175738 | 0.000164306 | 0.000187912 | 0.00018464 |

| | | | | | | | | | | | | | | | | | |
|---|---|---|---|---|---|---|---|---|---|---|---|---|---|---|---|---|---|
| 83 | alpha-Ionone | 127-41-3 | $C_{13}H_{20}O$ | 17.2 | 1874 | Terpene | Sweet, woody, floral, violet, orris, tropical, fruity | 0.002100297 | 0.001270786 | 0.001921168 | 0.001877712 | 0.001269374 | 0.002012284 | 0.001886774 | 0.001812874 | 0.001802384 | 0.001933268 |
| 84 | beta-Ionone epoxide | 23267-57-4 | $C_{13}H_{20}O_2$ | 18.9 | 2019 | Terpene | Fruity, sweet, berry, woody, violet, orris, powdery | 0.001456673 | 0.001330894 | 0.001314103 | 0.001370537 | 0.001495476 | 0.001394885 | 0.001381675 | 0.001193137 | 0.001325855 | 0.001481914 |
| 85 | Neophytadiene | 504-96-1 | $C_{20}H_{38}$ | 17.9 | 1929 | Terpene | - | 0.000179889 | 0.000251886 | 0.000341725 | 0.000260144 | 0.000364417 | 0.000273054 | 0.000324966 | 0.00032797 | 0.000322565 | 0.000437689 |
| 86 | trans-beta-Ionone | 79-77-6 | $C_{13}H_{20}O$ | 18.3 | 1966 | Terpene | dry powdery floral woody orris berry seedy | 0.001467838 | 0 | 0.004007879 | 0.001349576 | 0 | 0.004001712 | 0.003838057 | 0.003551018 | 0.003691388 | 0.003880322 |
| 87 | trans-Geranylacetone | 3796-70-1 | $C_{13}H_{22}O$ | 17.1 | 1865 | Terpene | Floral, fresh, green, rose | 0.000366719 | 0.000322123 | 0.000318789 | 0.000209751 | 0.000342013 | 0.000337794 | 0.000332665 | 0 | 0 | 0 |
| 88 | 2-Acetylthiazole | 24295-03-2 | $C_5H_5NOS$ | 14.3 | 1665 | Others | Pandan, popcorn, hazelnut, peanut | 4.19906E-05 | 5.78992E-05 | 6.20847E-05 | 3.29585E-05 | 7.25037E-05 | 6.0368E-05 | 1.98643E-05 | 1.23594E-06 | 0 | 0 |

| | | | | | | | | | | | | | | | | | |
|---|---|---|---|---|---|---|---|---|---|---|---|---|---|---|---|---|---|
| 89 | 2,3,5-Trimethylpyrazine | 14667-55-1 | C7H10N2 | 10.6 | 1421 | Pyrazine | Nutty | 5.56146E-06 | 6.45521E-06 | 1.25676E-05 | 0 | 0 | 0 | 4.92684E-06 | 1.19941E-05 | 1.3142E-05 | 1.35664E-05 |
| 90 | 2,5-Dimethylpyrazine | 123-32-0 | C6H8N2 | 9.3 | 1340 | Pyrazine | Cocoa, roasted, beefy, nutty | 2.83319E-05 | 0 | 0 | 2.30964E-05 | 1.68021E-05 | 0 | 1.85597E-05 | 0 | 4.28933E-05 | 1.95621E-05 |
| 91 | 2,6-dimethylpyrazine | 108-50-9 | C6H8N2 | 9.3 | 1338 | Pyrazine | Cocoa, roasted, nutty, meaty | 0 | 0 | 0 | 0 | 0 | 0 | 0 | 1.70375E-05 | 0 | 1.01784E-05 |
| 92 | 2-ethyl-3,6-dimethylpyrazine | 13360-65-1 | C8H12N2 | 11.2 | 1460 | Pyrazine | Roasted, potato | 2.4717E-05 | 2.25479E-05 | 2.13461E-05 | 1.89756E-05 | 2.06317E-05 | 2.2107E-05 | 5.59949E-06 | 2.93993E-05 | 1.99367E-05 | 1.892E-05 |
| 93 | 2-Ethyl-5-methylpyrazine | 13360-64-0 | C7H10N2 | 10.4 | 1408 | Pyrazine | Coffee, nut | 8.4125E-06 | 1.40265E-05 | 1.44929E-05 | 1.0381E-05 | 1.704E-05 | 1.73075E-05 | 1.08178E-05 | 8.73359E-06 | 1.25065E-05 | 1.31454E-05 |
| 94 | 2-Ethylpyrazine | 13925-00-3 | C6H8N2 | 9.5 | 1350 | Pyrazine | Nutty, fermented coffee, meaty | 0 | 1.4925E-06 | 0 | 0 | 0 | 0 | 0 | 0 | 0 | 0 |
| 95 | 2-Methylpyrazine | 109-08-0 | C5H6N2 | 8.4 | 1281 | Pyrazine | Nutty, cocoa, chocolate | 5.64735E-06 | 1.56795E-05 | 1.67164E-05 | 1.50864E-05 | 1.66667E-05 | 1.35332E-05 | 0 | 1.45025E-05 | 1.58873E-05 | 1.60826E-05 |

| | | | | | | | | | | | | | | | |
|---|---|---|---|---|---|---|---|---|---|---|---|---|---|---|---|
| 96 | 2,4,6-trimethylpyridine | 108-75-8 | C8H11N | 10.0 | 1388 | Pyridine | - | 5.35797E-06 | 6.96537E-06 | 8.23105E-06 | 6.81916E-06 | 7.76736E-06 | 8.21101E-06 | 4.28681E-06 | 7.05771E-06 | 4.94132E-06 | 5.82558E-06 |
| 97 | 3-methylpyridine | 108-99-6 | C6H7N | 8.9 | 1316 | Pyridine | Green, earthy, hazelnut, nutty | 4.72888E-06 | 3.89947E-06 | 0 | 7.09646E-06 | 0 | 3.85177E-06 | 0 | 0 | 8.36633E-06 | 4.07514E-06 |
| 98 | Pyridine | 110-86-1 | C5H5N | 7.2 | 1204 | Pyridine | Sour, fishy | 1.01316E-05 | 9.70381E-06 | 1.02752E-05 | 7.86009E-06 | 6.04669E-06 | 1.05357E-05 | 1.15231E-05 | 1.07348E-05 | 9.52469E-06 | 9.12224E-06 |
| 99 | 1H-Pyrrole-2,5-dione, 3-ethyl-4-methyl- | 21494-57-5 | C7H9NO | 21.2 | 2298 | Pyrrole | - | 0.000192216 | 0.000163596 | 0.000182589 | 0.000195079 | 0.000218631 | 0.000170895 | 0.00018866 | 0.00016829 | 0.000178433 | 0.000196725 |
| 101 | Dimethyl trisulfide | 3658-80-8 | C2H6S3 | 10.2 | 1399 | Sulphurous | Sulphurous, onion, meaty, rotten eggs | 0 | 0 | 0 | 2.53319E-06 | 0 | 0 | 0 | 0 | 0 | 0 |

Summary of all the volatile compounds detected by GC-MS in unfermented microalgae samples and *L. plantarum*-fermented microalgae samples across different time points (0h, 24h, 48h, 72h, and 96h). Each compound entry includes its chemical name, CAS number, molecular formula, retention time (min), retention index (RI), compound class, and odour description and abundances at different time points.

##### Table S4: ASTREE E-tongue sensor summary with food applications

| | Sensor | Taste Modality | Sensitivity and typical detected compounds | Common applications |
|---|---|---|---|---|
| 1 | AHS | Sour | Organic acids, $H^+$ ions | · Fermented foods (e.g., yogurt, kimchi, sauerkraut).<br>· Vinegar/acidity profiling in sauces.<br>· Spoilage detection (e.g., acid build-up). |
| 2 | CTS | Salty | $Na^+$, $K^+$, $Cl^-$, mineral salts | · Salt reduction in soups and snacks.<br>· Mineral water profiling.<br>· Electrolyte beverage development. |
| 3 | NMS | Umami | Glutamate, IMP, GMP, amino acids | · Umami enhancement in broth/stock.<br>· Fermented soy (miso, soy sauce).<br>· Meat analogues or plant-based meat optimization. |
| 4 | ANS | Sweet | Glucose, fructose, sucrose, and artificial sweeteners | · Sweetener formulation in beverages<br>· Sugar content consistency in fruit juices<br>· Sugar-free product comparisons |
| 5 | SCS | Bitter | Caffeine, polyphenols, quinine, alkaloids | · Coffee and tea bitterness profiling.<br>· Cocoa/chocolate evaluation.<br>· Bitterness masking in functional foods/supplements. |
| 6 | PKS | General/Reference | Detects overall ionic strength or changes in general chemical environment | · Acts as a reference to help differentiate complex mixtures and stabilize the signal |
| 7 | CPS | General/Reference | | |

Description of electronic tongue (E-tongue) sensors used for taste modality detection in food samples. The table summarizes the target analytes for each sensor and their common applications in food quality evaluation, flavour profiling, and product development [5-13].

##### Table S5. Metabolite profile of umami-related compounds detected during *L. plantarum*-mediated biotransformation of microalgae

| Compound name | Mass to charge ratio (m/z) | Peak Area | | | | |
|---|---|---|---|---|---|---|
| | | ***L. plantarum*_MA_0h** | ***L. plantarum*_MA_24h** | ***L. plantarum*_MA_48h** | ***L. plantarum*_MA_72h** | ***L. plantarum*_MA_96h** |
| Glutamic acid | 148.0605 | 26077.91 | 25747.33 | 7396.46 | 6104.25 | 4747.83 |

| | | | | | | |
|---|---|---|---|---|---|---|
| 3'-AMP | 346.0547 | 15341.44 | 15621.17 | 64523.82 | 71823.88 | 56305.31 |
| Adenosine 2',3'-cyclic monophosphate (cAMP) | 330.059 | 10029.51 | 8643.28 | 14599.89 | 8634.29 | 4084.71 |
| Adenosine 5'-monophosphate (AMP) | 348.07 | 1293.7 | 13687.83 | 57535.56 | 68646.83 | 57002.32 |
| Cyclic GMP (cGMP) | 344.0396 | 4024.884766 | 4647.526367 | 7276.584961 | 3959.945313 | 774.7654419 |
| Guanosine 5'-monophosphate | 362.0501 | 40930.2 | 48802.02 | 61079.58 | 32945.93 | 15488.41 |

Umami-related compounds were detected using LC-MS and identified based on their mass-to-charge ratio (m/z) Peak area values represent relative abundances at each fermentation time point (0h, 24h, 48h, 72h, 96h).

**Table S6. Metabolite profile of amino acids and their bioconversion products during *L. plantarum*-mediated biotransformation of microalgae**

| **Compound name** | **Mass to charge ratio (m/z)** | **Peak Area** | | | | |
|---|---|---|---|---|---|---|
| | | ***L. plantarum*_MA_0h** | ***L. plantarum*_MA_24h** | ***L. plantarum*_MA_48h** | ***L. plantarum*_MA_72h** | ***L. plantarum*_MA_96h** |
| L-Tyrosine | 182.0787 | 37898 | 34893 | 7181 | 0 | 0 |
| Phenylalanine | 166.0866 | 89252 | 89795 | 8013 | 6714 | 4542 |
| 4-Hydroxyphenyllactic acid | 181.0503 | 7257 | 5588 | 87693 | 127266 | 145663 |
| D-Tryptophan | 205.0973 | 58778 | 57684 | 8754 | 16172 | 13113 |
| D-3-phenyllactic acid | 165.0557 | 28351 | 20197 | 136790 | 327943 | 389709 |
| DL-Indole-3-lactic acid | 204.0666 | 0 | 0 | 24333.55 | 53092.48 | 60450.36 |

Amino acids and their relevant metabolites were detected using LC-MS and identified based on their mass-to-charge ratio (m/z). Peak area values represent relative abundances at each fermentation time point (0h, 24h, 48h, 72h, 96h).

**Table S7. Demographic and lifestyle information of the sniff test panellists (n = 10).**

| Participant (n=10) | Gender | Age range | Race | Diet | Smoker? | Self-rated smell sensitivity |
|---|---|---|---|---|---|---|
| 1 | Male | 31-40 | Chinese | Omnivore | No | 5 |
| 2 | Male | 41-50 | Chinese | Omnivore | No | 2 |
| 3 | Male | 31-40 | Chinese | Omnivore | No | 3 |
| 4 | Male | 21-30 | Chinese | Omnivore | No | 3 |
| 5 | Male | 31-40 | Chinese | Omnivore | No | 3 |
| 6 | Female | 31-40 | Chinese | Omnivore | No | 5 |
| 7 | Female | 41-50 | Indian | Omnivore | No | 3 |
| 8 | Female | 31-40 | Malay | Omnivore | No | 5 |
| 9 | Female | 21-30 | Chinese | Omnivore | No | 3 |
| 10 | Female | 31-40 | Chinese | Omnivore | No | 3 |

Demographic details and lifestyle factors of the untrained olfactory evaluation panel (n = 10). All panellists were non-smokers and followed an omnivorous diet. Self-rated olfactory sensitivity was collected using a 5-point scale (1 = very poor, 5 = very strong).

**Table S8. Two-way ANOVA statistics for grassy odour intensity.**

| **ANOVA table** | **SS** | **DF** | **MS** | **F (DFn, DFd)** | **P value** | **Significance and Interpretation** |
|---|---|---|---|---|---|---|
| Interaction (Time × Treatment) | 3.400 | 4 | 0.8500 | F (4, 72) = 3.161 | P=0.0188 | The change in grassy odour over time depends on treatment. |
| Time (Row Factor) | 2.440 | 4 | 0.6100 | F (3.282, 59.07) = 2.269 | P=0.0845 | No significant change in grassy odour intensity over time alone. |
| Treatment (Column Factor) | 0.2500 | 1 | 0.2500 | F (1, 18) = 0.2089 | P=0.6531 | No overall difference between fermented and unfermented samples. |
| Subject (Panellist variation) | 21.54 | 18 | 1.197 | F (18, 72) = 4.450 | P<0.0001 | Panellists varied significantly in how they rated the grassy odour. |

A two-way repeated measures ANOVA was used to assess the effects of fermentation treatment, time, and their interaction on grassy odour intensity. A significant interaction between time and treatment ($p = 0.0188$) indicates that the change in grassy odour perception over time differed depending on fermentation condition. A significant subject effect indicates variability among panellist ratings ($P<0.0001$). No significant main effects of time or treatment were observed. Geisser-Greenhouse correction was applied to the time effect ($\varepsilon = 0.8204$). Note: $F$ = F-ratio; DF = degrees of freedom; SS = sum of squares; MS = mean square.

**Table S9. Statistical summary of pairwise comparison for grassy odour intensity between unfermented and *L. plantarum*-fermented microalgae.**

| Difference between column means | |
|---|---|
| Mean of Microalgae only | 2.060 |
| Mean of *L. plantarum*_MA | 1.960 |
| Difference between means | 0.1000 |
| SE of difference | 0.2188 |
| 95% CI of difference | -0.3596 to 0.5596 |

Reported values include mean difference, standard error (SE), and 95% confidence interval (CI).

**Table S10. Two-way ANOVA statistics for sour odour intensity.**

| ANOVA table | SS | DF | MS | F (DFn, DFd) | P value | Significance and Interpretation |
|---|---|---|---|---|---|---|
| Interaction (Time × Treatment) | 1.840 | 4 | 0.4600 | F (4, 72) = 2.875 | P=0.0287 | The change in sour odour over time depends on treatment. |
| Time (Row Factor) | 1.840 | 4 | 0.4600 | F (2.301, 41.41) = 2.875 | P=0.0607 | No significant change in sour odour intensity over time alone. |
| Treatment (Column Factor) | 0.8100 | 1 | 0.8100 | F (1, 18) = 0.9480 | P=0.3431 | No overall difference in sour odour between the fermented and unfermented groups. |
| Subject (Panellist variation) | 15.38 | 18 | 0.8544 | F (18, 72) = 5.340 | P<0.0001 | Panellists varied significantly in how they rated sour odour. |

A two-way repeated measures ANOVA was used to assess the effects of fermentation treatment, time, and their interaction on sour odour intensity. A significant interaction between time and treatment (p = 0.0287) indicates that the change in grassy odour perception over time differed depending on fermentation condition. A significant subject effect indicates variability among panellist ratings (P<0.0001). No significant main effects of time or treatment were observed. Geisser-Greenhouse correction was applied to the time effect (ε = 0.5751). Note: *F* = F-ratio; DF = degrees of freedom; SS = sum of squares; MS = mean square.

**Table S11. Statistical summary of pairwise comparison for sour odour intensity between unfermented and *L. plantarum*-fermented microalgae.**

| **Difference between column means** | |
|---|---|
| Mean of Microalgae only | 1.220 |
| Mean of *L. plantarum*_MA | 1.400 |
| Difference between means | -0.1800 |
| SE of difference | 0.1849 |
| 95% CI of difference | -0.5684 to 0.2084 |

Reported values include mean difference, standard error (SE), and 95% confidence interval (CI).

**Table S12. Two-way ANOVA statistics for fishy odour intensity.**

| **ANOVA table** | **SS** | **DF** | **MS** | **F (DFn, DFd)** | **P value** | **Significance and Interpretation** |
|---|---|---|---|---|---|---|
| Interaction (Time × Treatment) | 0.1000 | 4 | 0.02500 | F (4, 72) = 0.06329 | P=0.9925 | No difference in changes between fermented and unfermented samples over time. |
| Time | 1.460 | 4 | 0.3650 | F (2.627, 47.29) = 0.9241 | P=0.4262 | No significant change in fishy |

| | | | | | | |
|---|---|---|---|---|---|---|
| (Row Factor) | | | | | | odour intensity over time alone. |
| Treatment (Column Factor) | 0.000 | 1 | 0.000 | F (1, 18) = 0.000 | P>0.9999 | No overall difference in fishy odour between the fermented and unfermented groups. |
| Subject (Panelist variation) | 24.96 | 18 | 1.387 | F (18, 72) = 3.511 | P<0.0001 | Panellists varied significantly in how they rated fishy odour. |

A two-way repeated measures ANOVA was used to assess the effects of fermentation treatment, time, and their interaction on fishy odour intensity. A significant subject effect indicates variability among panellist ratings (P<0.0001). No significant main effects of interaction, time or treatment were observed. Geisser-Greenhouse correction was applied to the time effect ($\varepsilon = 0.6569$). Note: *F* = F-ratio; DF = degrees of freedom; SS = sum of squares; MS = mean square.

**Table S13. Statistical summary of pairwise comparison for fishy odour intensity between unfermented and *L. plantarum*-fermented microalgae.**

| **Difference between column means** | |
|---|---|
| Mean of Microalgae only | 1.520 |
| Mean of *L. plantarum*_MA | 1.520 |
| Difference between means | 0.000 |
| SE of difference | 0.2355 |
| 95% CI of difference | 1.520 |

Reported values include mean difference, standard error (SE), and 95% confidence interval (CI).

**Table S14. Two-way ANOVA statistics for earthy odour intensity.**

| ANOVA table | SS | DF | MS | F (DFn, DFd) | P value | Significance and Interpretation |
|---|---|---|---|---|---|---|
| Interaction (Time × Treatment) | 0.9400 | 4 | 0.2350 | F (4, 72) = 0.7719 | 0.9400 | No difference in changes between fermented and unfermented samples over time. |
| Time (Row Factor) | 1.140 | 4 | 0.2850 | F (3.299, 59.38) = 0.9361 | 1.140 | No significant change in earthy odour intensity over time alone. |
| Treatment (Column Factor) | 0.3600 | 1 | 0.3600 | F (1, 18) = 0.1617 | 0.3600 | No overall difference in earthy odour between the fermented and unfermented groups. |
| Subject (Panellist variation) | 40.08 | 18 | 2.227 | F (18, 72) = 7.314 | 40.08 | Panellists varied significantly in how they rated earthy odour. |

A two-way repeated measures ANOVA was used to assess the effects of fermentation treatment, time, and their interaction on earthy odour intensity. A significant subject effect indicates variability among panellist ratings ($P<0.0001$). No significant main effects of interaction, time or treatment were observed. Geisser-Greenhouse correction was applied to the time effect ($\varepsilon = 0.5470$). Note: $F$ = F-ratio; DF = degrees of freedom; SS = sum of squares; MS = mean square.

**Table S15. Statistical summary of pairwise comparison for earthy odour intensity between unfermented and *L. plantarum*-fermented microalgae.**

| **Difference between column means** | |
|---|---|
| Mean of Microalgae only | 1.720 |

| | |
|---|---|
| Mean of *L. plantarum*_MA | 1.600 |
| Difference between means | 0.1200 |
| SE of difference | 0.2984 |
| 95% CI of difference | -0.5070 to 0.7470 |

Reported values include mean difference, standard error (SE), and 95% confidence interval (CI).

#### Supporting Figures

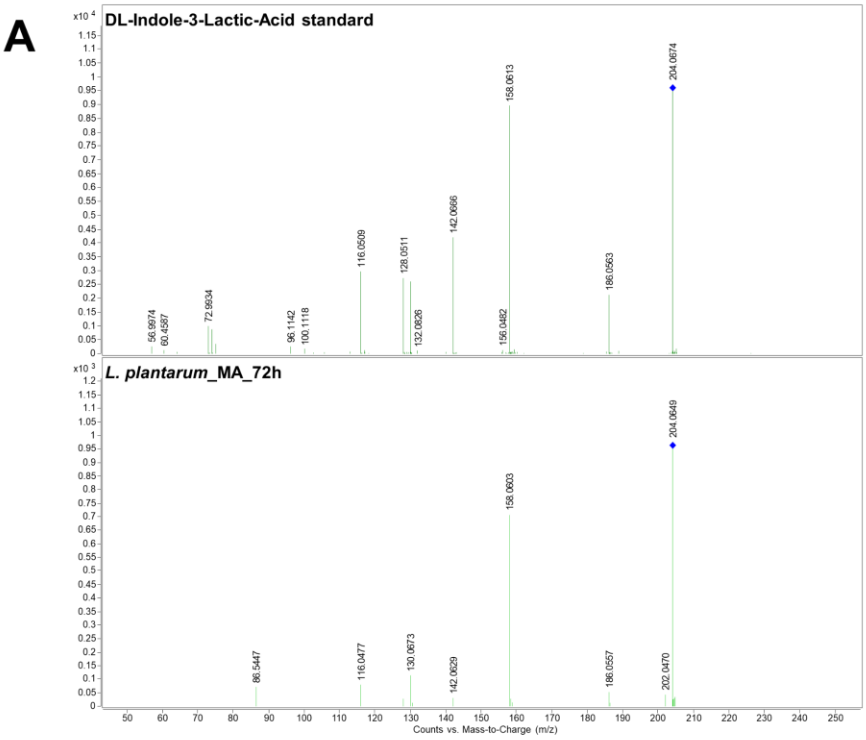

B

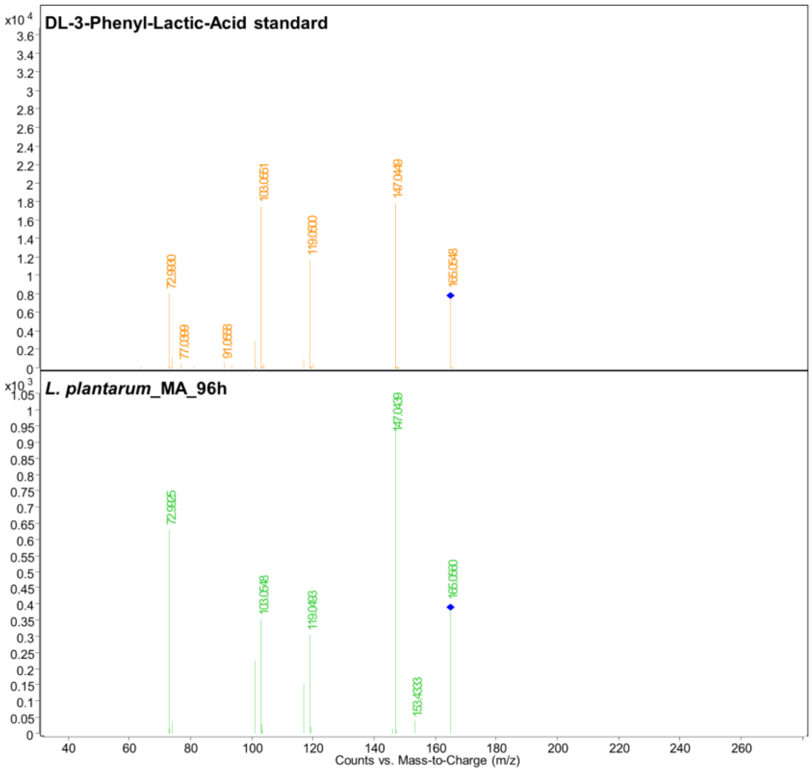
DL-3-Phenyl-Lactic-Acid standard
x10 4
72.9930
77.0399
91.0558
103.0551
119.0500
147.0449
165.0548
L. plantarum_MA_96h
x10 3
72.9925
103.0548
119.0483
147.0439
153.4333
165.0560
Counts vs. Mass-to-Charge (m/z)

C

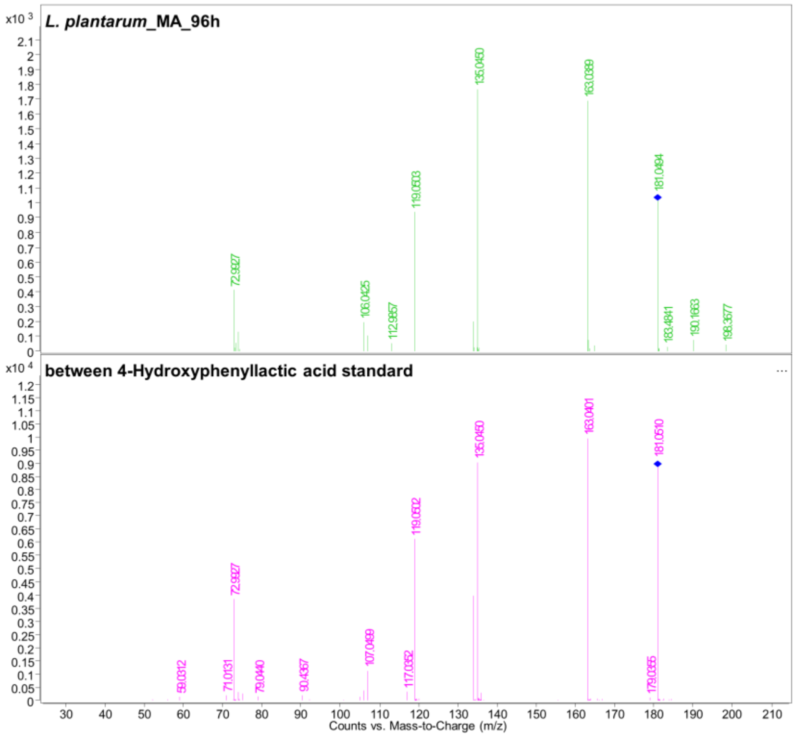
L. plantarum_MA_96h
x10 3
72.9927
106.0425
112.9857
119.0503
135.0450
163.0389
181.0494
183.4841
190.1663
198.3577
between 4-Hydroxyphenyllactic acid standard
x10 4
59.0312
71.0131
72.9927
79.0440
90.4367
107.0499
117.0352
119.0502
135.0450
163.0401
179.0355
181.0510
Counts vs. Mass-to-Charge (m/z)

**Figure S1. LC-MS/MS fragmentation spectra at collision-induced dissociation (CID)10V of aromatic lactic acid derivatives in *L. plantarum*-fermented microalgae. (A)** MS/MS spectra of key fragment ions matched between DL-Indole-3-lactic acid reference standard and *L. plantarum*-fermented microalgae at 72h of fermentation. **(B)** MS/MS spectra of key fragment ions matched between DL-3-phenyllactic acid reference standard and *L. plantarum*-fermented microalgae at 96 h of fermentation. **(C)** MS/MS spectra of key fragment ions matched between 4-Hydroxyphenyllactic acid reference standard and *L. plantarum*-fermented microalgae at 96h of fermentation.

A

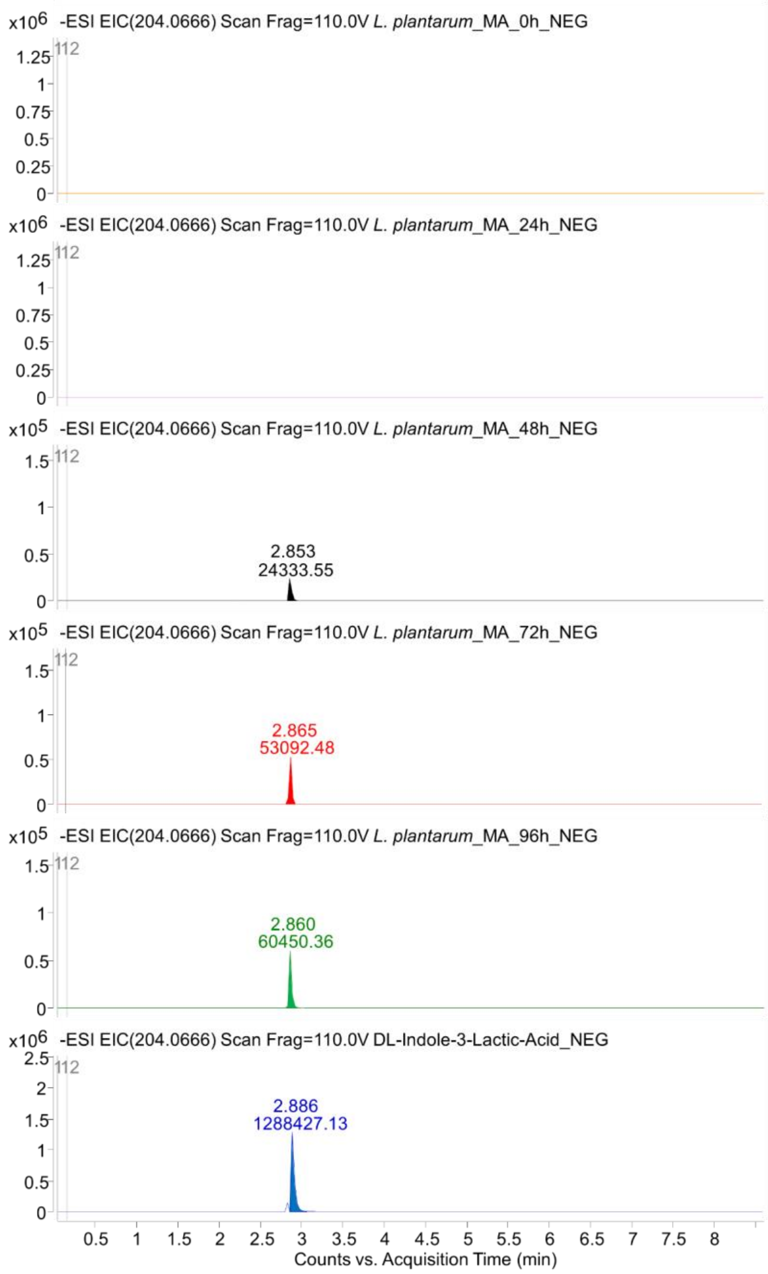

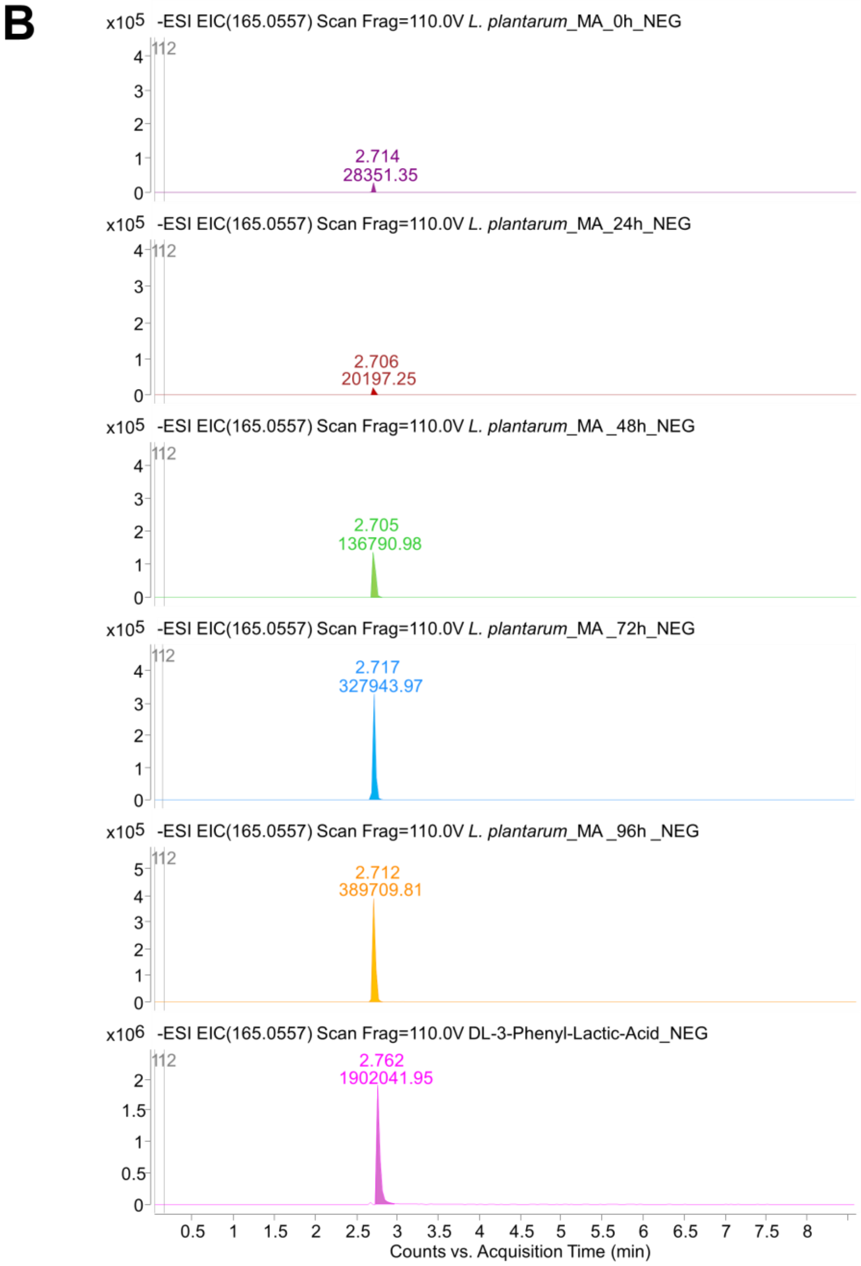
B
x10^5 -ESI EIC(165.0557) Scan Frag=110.0V L. plantarum_MA_0h_NEG
2.714
28351.35
x10^5 -ESI EIC(165.0557) Scan Frag=110.0V L. plantarum_MA_24h_NEG
2.706
20197.25
x10^5 -ESI EIC(165.0557) Scan Frag=110.0V L. plantarum_MA _48h_NEG
2.705
136790.98
x10^5 -ESI EIC(165.0557) Scan Frag=110.0V L. plantarum_MA _72h_NEG
2.717
327943.97
x10^5 -ESI EIC(165.0557) Scan Frag=110.0V L. plantarum_MA _96h _NEG
2.712
389709.81
x10^6 -ESI EIC(165.0557) Scan Frag=110.0V DL-3-Phenyl-Lactic-Acid_NEG
2.762
1902041.95
Counts vs. Acquisition Time (min)

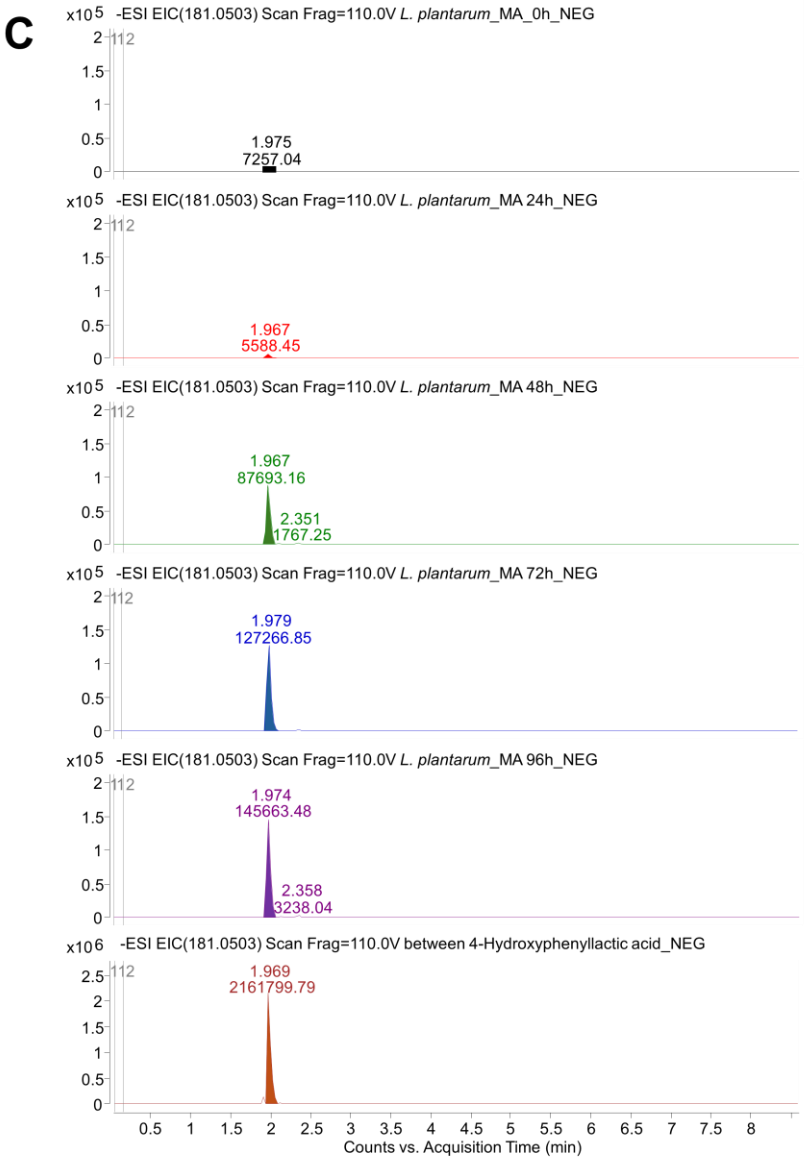

**Figure S2. Extracted ion chromatograms (EICs) from LC-MS showing the bioconversion of precursor amino acids during *L. plantarum*-mediated fermentation of microalgae (MA) over 96 hours. (A–C)** LC-MS EICs showing the time-dependent increase in signal intensity of **(A)** DL-indole-3-lactic acid (m/z 204.0666), **(B)** D-3-phenyllactic acid (m/z 165.0557), and **(C)** 4-Hydroxyphenyllactic acid (m/z 181.0503), across fermentation time points (0h, 24h, 48h, 72h, 96h) compared to reference standards.

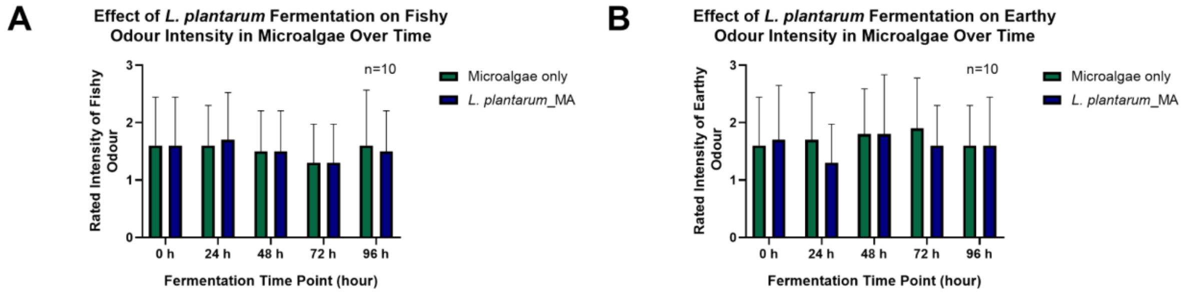

**Figure S3. Olfactory evaluation of microalgae fermented with *L. plantarum.* (A-B)** Sensory intensities of **(A)** fishy and **(B)** earthy odour attributes in unfermented microalgae (Microalgae only) and *L. plantarum*-fermented microalgae (*L. plantarum*_MA) over 96 hours. Olfactory evaluation was performed by an untrained in-house olfactory panel (n = 10), with intensities ranked on a structured 3-point scale (1 = weak, 2 = moderate, 3 = strong). Bars represent mean ± standard deviation.

**Supporting References**

1 X. H. Chin, R. Soh, G. Chan, P. Ng, A. Thong, H. Elhalis, K. Yoganathan, Y. Chow and S. Q. Liu, *Current Research in Food Science*, 2025, **10**, 100933.
2 F. L. Sirota, F. Goh, K.-N. Low, L.-K. Yang, S. C. Crasta, B. Eisenhaber, F. Eisenhaber, Y. Kanagasundaram and S. B. Ng, *Journal of Genomics*, 2018, **6**, 63–73.
3 R. Schmid, S. Heuckeroth, A. Korf, A. Smirnov, O. Myers, T. S. Dyrlund, R. Bushuiev, K. J. Murray, N. Hoffmann, M. Lu, A. Sarvepalli, Z. Zhang, M. Fleischauer, K. Dührkop, M. Wesner, S. J. Hoogstra, E. Rudt, O. Mokshyna, C. Brungs, K. Ponomarov, L. Mutabdžija, T. Damiani, C. J. Pudney, M. Earll, P. O. Helmer, T. R. Fallon, T. Schulze, A. Rivas-Ubach, A. Bilbao, H. Richter, L.-F. Nothias, M. Wang, M. Orešič, J.-K. Weng, S. Böcker, A. Jeibmann, H. Hayen, U. Karst, P. C. Dorrestein, D. Petras, X. Du and T. Pluskal, *Nature Biotechnology*, 2023, **41**, 447–449.
4 Z. Pang, Y. Lu, G. Zhou, F. Hui, L. Xu, C. Viau, A. F. Spigelman, P. E. MacDonald, D. S. Wishart, S. Li and J. Xia, *Nucleic Acids Research*, 2024, **52**, 398–406.
5 B. Zhou, X. Zhao, L. Laghi, X. Jiang, J. Tang, X. Du, C. Zhu and G. Picone, *Foods*, 2024, **13**, 2911.
6 L. Wang, J. Xie, Q. Wang, J. Hu, Y. Jiang, J. Wang, H. Tong, H. Yuan and Y. Yang, *Food Chemistry: X*, 2024, **23**, 101519.
7 M. Gil, M. Rudy, P. Duma-Kocan and R. Stanisławczyk, *Applied Sciences*, 2025, **15**, 1530.
8 M. S. Moazzem, M. Hayden, D.-J. Kim and S. Cho, *Foods*, 2024, **13**, 3269.
9 N. Muenprasitivej, Electronic Tongue Analysis of Major and Minor Steviol Glycosides and Their Application in Foods, https://etd.auburn.edu/handle/10415/8305.
10 R. I. Potter, J. Lee and C. F. Ross, *Food science & nutrition*, 2025, **13**, e70366.
11 T. Wang, L. Yang, Y. Xiong, B. Wu, Y. Liu, M. Qiao, C. Zhu, H. Wu, J. Deng and J. Guan, *Frontiers in Nutrition*, 2024, **11**, 1435364.
12 X. Cai, Y. Zeng, K. Zhu, Y. Peng, P. Xv, P. Dong, M. Qiao and W. Fan, *Foods*, 2025, **14**, 1982.
13 X. Yin, Y. Lv, R. Wen, Y. Wang, Q. Chen and B. Kong, *Meat Science*, 2021, **172**, 108345.
